# Differential Reactivation of Task-Demand-Associated Firing Patterns in Subicular and CA1 Place Cells during a Hippocampal Memory Task

**DOI:** 10.1101/2024.07.24.605041

**Authors:** Jae-Min Seol, Su-Min Lee, Inah Lee

## Abstract

Reactivation of place cells during sharp-wave ripples (SWRs) in the hippocampus is pivotal for memory consolidation, yet differences in SWR dynamics between the hippocampus and its neighboring subiculum remain underexplored. We examined the differential reactivations of task-demand-associated representations during SWR events in the subiculum and CA1 during a visual scene memory task in rats. In the task, the spiking activity of place cell ensembles was reactivated during a SWR event according to task demands. These reactivations were more frequent and were associated with more heterogeneous task-demand types in the subiculum compared with the CA1. Neural manifold analysis showed that the neural states of the reactivated ensemble were more clearly clustered into distinct states during subicular SWRs according to the task-demand-associated variables. These subicular characteristics were driven by multiple subfields of the subicular place field, parcellated by the theta phase precession cycle. In contrast, CA1 exhibited a higher incidence of spatial replay than the subiculum. These findings indicate that the subiculum plays a key role in transmitting task-specific variables from the hippocampus to other brain regions.

## Introduction

Episodic memory involves the recollection of events within a spatial context. Such memories comprise not only the locations where events occurred but also encompass other related elements, such as objects and background contexts. It has been widely postulated that the hippocampus is vital for both the formation and retrieval of these event memories (*1,2*). Additionally, research indicates that the consolidation of event memory requires specific physiological mechanisms in the hippocampus, particularly sharp wave ripples (SWRs), which are believed to play a crucial role during memory consolidation in learning (3–5).

The literature suggests that SWRs facilitate the reinstatement of spatial information in the hippocampus through the ‘replay’ of place cells according to their spatial firing patterns observed during prior exploration (*6–8*). Numerous studies have demonstrated that task-related information can influence these replay patterns. For instance, the amount of reward (*9–11*) or the valence associated with a location (*12–14*) modulates neural activity during SWRs by affecting the participation rates of certain cell types or directing robust replays toward significant locations. Moreover, replay sequences are responsive to obstacles or shortcuts (*15,16*), and place cells’ firing rates are adjusted based on contextual differences (*17–19*). These findings indicate that task-relevant factors influence the spatial reactivation patterns of place cells. However, given that the firing patterns of place cells are modulated by task-related information, there is limited understanding of how such task-related spiking activities of place cells manifest during SWRs apart from the spatial replay.

It is well established that SWR-dependent reactivation of neural activity is a robust phenomenon in the hippocampus that is critical for learning and memory (*4,5,20–25*). However, the role of the subiculum – the interface between the hippocampus and cortical regions – in this process remains largely unexplored. The subiculum has been proposed as a crucial pathway for transmitting hippocampal activity, including SWRs, to neocortical areas (*26*), or as a region capable of generating SWRs independently (*27*). It has also been reported that firing rates of a subset of subicular neurons either increase or decrease during SWRs (*28,29*). Nonetheless, it is unclear whether subicular neurons are reactivated in relation to the spatial and non-spatial information of event memory during SWRs. The cells in the subiculum is known for their spatially broadly tuned firing patterns compared with CA1 place cells (*30–34*). Such differences in firing patterns between the CA1 and subiculum also suggest potential variations in reactivation patterns during SWRs between the two regions.

Our previous research (*32,34*) identified distinct firing properties of subicular neurons for representing task-relevant information in a hippocampal memory task. Specifically, in a visual scene memory (VSM) task, place cells in both the subiculum and CA1 demonstrated firing-rate modulations (rate remapping) based on visual scenes in the background or on the impending arm choice. These rate-remapping events were observed at the level of the θ-phase–based subfields defined by the θ-phase precession of spiking phases. Additionally, the θ-phase–based subfields of subicular neurons showed stronger modulation of firing rates in response to critical task variables (i.e., visual scenes and associated choice arms), thus termed task-demand–associated (TDA) variables in this study. Building on our previous findings, we hypothesize that subicular neuronal ensembles are coactivated during SWRs to represent TDA information more prominently than hippocampal ensembles.

## Results

### Behavioral Performance in a Hippocampal-Dependent VSM Task

The data utilized in this study were originally collected in our previous research (*32*). Five rats were trained to associate four distinct visual scene stimuli with either the left or right arm of a T-maze as part of a VSM task (**Fig. 1**). At the beginning of each trial, the guillotine door of the start box was opened, allowing the rat to walk onto the maze stem, while one of the four visual scenes – zebra, pebbles, bamboo, and mountain (pseudo-randomly selected for each trial) – was displayed on an array of three LCD monitors surrounding the choice area of the maze. The task required the rat to visit the arm associated with the presented scene stimulus to retrieve a cereal reward from the food well. After making a choice and returning to the start box with the cereal reward, the guillotine door was closed, signaling the start of the inter-trial interval (ITI). Our previous research indicated that the VSM task is dependent on the hippocampus (*35*) as well as the subiculum (*36*).

**Fig. 1.**
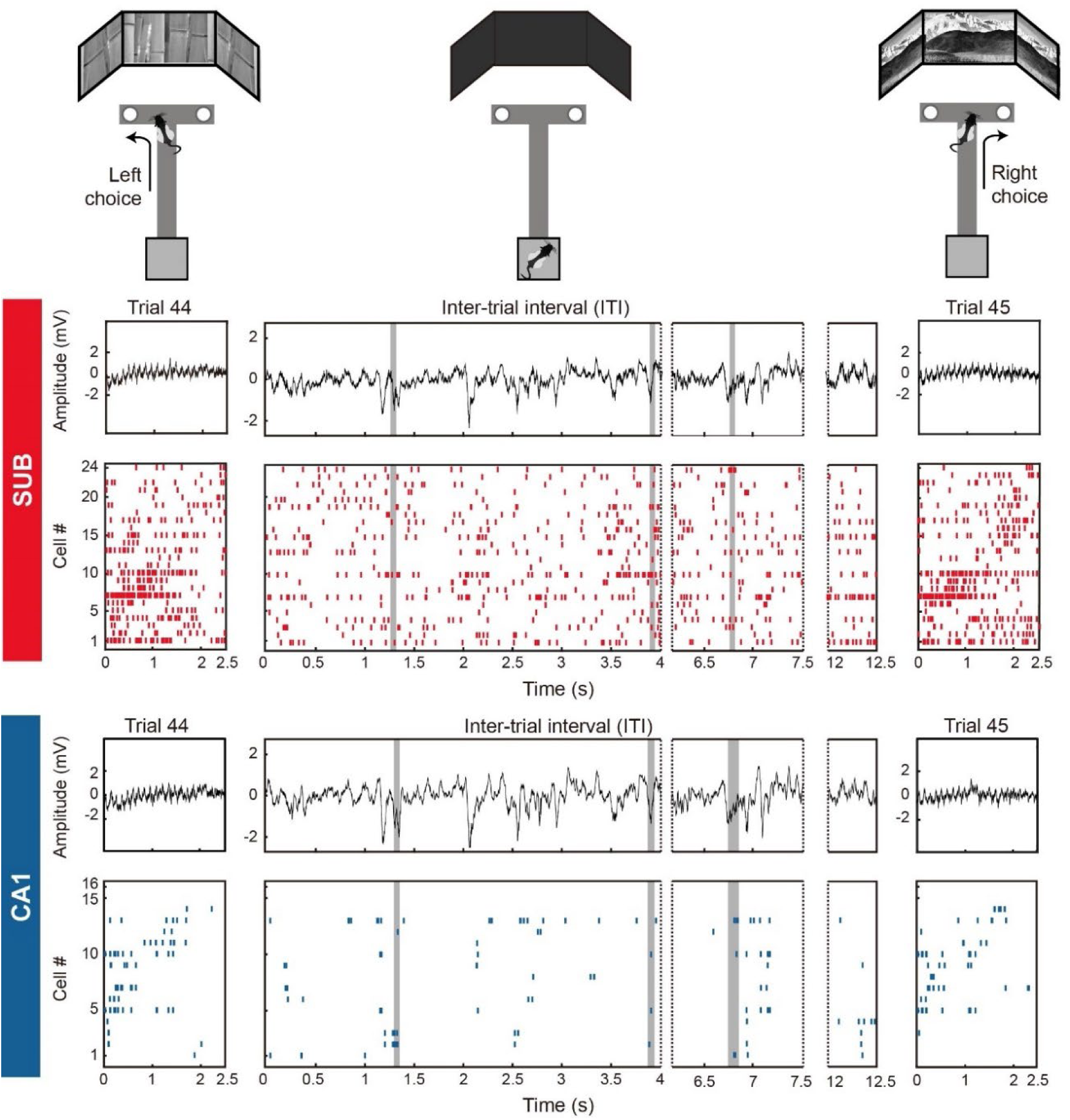
SWR detection in the subiculum and CA1 during the VSM task. Top: Illustration of trial epochs and inter-trial intervals in the VSM task. During each trial epoch, the rat was required to choose the arm (left or right) associated with the visual scene presented on the monitors at the end of the stem to obtain a cereal reward from the food well at the end of the arm. Once the rat returned to the start box, the guillotine door was closed, and the rat was required to stay inside the start box for ∼10 seconds. Bottom: LFP and spiking activity of cell ensembles recorded simultaneously in the subiculum and CA1 during an example ITI and the preceding and subsequent trials (trials 44 and 45, from rat 232-07 session). In the center column, SWR events are indicated by gray shading. In each region, cells are sorted by on-track field peak positions. SUB, subiculum.

By the time the physiological recording sessions began, all rats had surpassed the performance criterion (75% correct choices) for all scene stimuli (Z = 3.94, *p* < 0.001 for zebra; Z = 3.91, *p* < 0.001 for pebbles; Z = 3.75, *p* < 0.001 for bamboo; Z = 3.91, *p* < 0.001 for mountain; one-sample Wilcoxon signed rank test). There were no significant differences in ITIs across the different scene conditions (F(_3_) = 0.14, *p* = 0.936, one-way ANOVA; **Fig. S1**). We recorded spiking activity from single units and local field potentials (LFPs) simultaneously from the dorsal CA1 and subiculum while the rats performed the VSM task. Consistent with our previous findings (*32,34*), the average firing rate of complex-spiking cells (n = 192 in the subiculum; n = 282 in the CA1) was higher in the subiculum than in the CA1 (Z = 8.45, *p* < 0.001), whereas the spatial information score (bit/spike) was lower in the subiculum than in CA1 (Z = -12.51, *p* < 0.001; Wilcoxon rank-sum test).

SWRs were defined as LFP signals with high frequency (150-250 Hz) and large amplitude (>2 standard deviations [SDs] for the SWR peak and > 0.5 SD for SWR boundaries; see Material and Methods for details) (**Fig. 1**). SWRs analyzed in this study occurred during ITIs following correct trials when the rats were awake but immobile. We included a SWR event in further analyses only if three or more putative complex-spiking cells fired simultaneously (n = 5899 in the subiculum; n = 1899 in the CA1) (**Fig. 1**). SWRs in the subiculum were shorter in duration (Z = -3.11, *p* = 0.002; **Fig. S2**) and involved fewer coactivated cells per SWR (Z = -33.47, *p* < 0.001) compared with CA1 SWRs. Additionally, the amplitude of the filtered LFP was larger in the subiculum than in CA1 (Z = 31.74, *p* < 0.001; Wilcoxon rank-sum test).

### Selective Firing of Subiculum and CA1 Place Cells for Visual Scene Stimuli and Upcoming Choices

In our previous study (*34*), we demonstrated that the traditional spatial firing field (place field), defined by the firing rate (rate-based field’), could be further subdivided into multiple subfields (‘θ-phase-based fields’) based on the cell’s theta phase precession cycle. This method revealed spatially localized subfields within the broad firing field of a subicular neuron, making most firing properties of the subicular θ-phase–based fields comparable to those of rate-based fields in CA1. In the current study, we applied the same θ-phase–based field identification method to define θ-phase–based place fields in both the subiculum and CA1 (**Fig. 2A**).

**Fig. 2.**
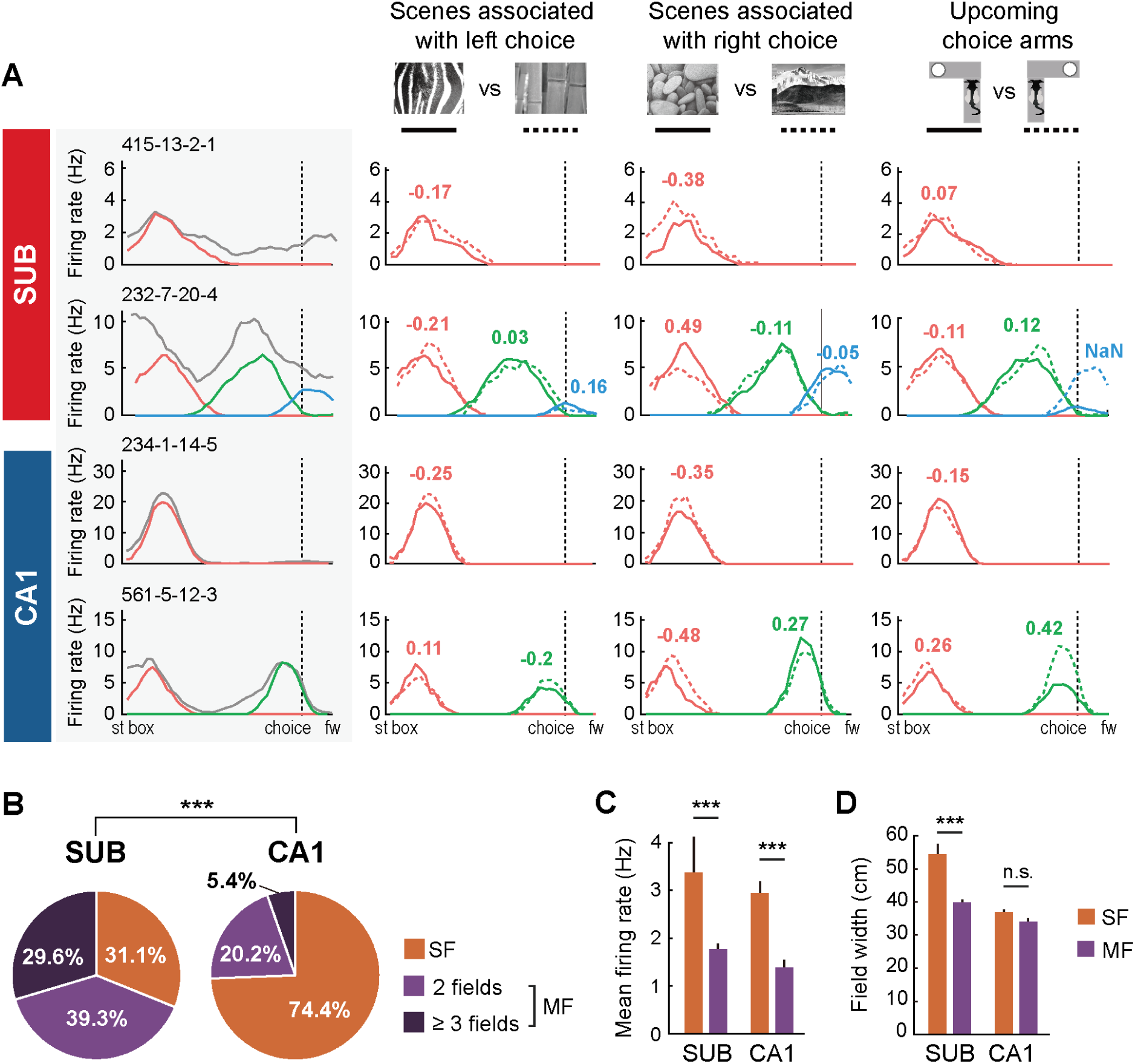
Rate modulation of θ-phase–based fields by scene and choice. **(A)** Each row displays an example cell from the subiculum and CA1. In the first column of each row (gray shaded) is the linearized firing rate map on the T-maze, averaged over all trials. Overall firing activity is shown as gray lines, whereas place fields defined by theta phases are shown as colored lines. The subsequent columns on the right depict firing rate maps constructed from trials associated with different TDA information (visual scenes and upcoming choice arms). In each column, solid and dotted lines represent the average firing patterns for different scenes or choice direction, respectively, color-coded by θ-phase–based subfields. The numbers above the fields indicate the selectivity indices for the scene pairs or the upcoming choice direction (see Materials and Methods for detailed description), and the number is shown in color when the absolute value of the selectivity index is > 0.1. **(B)** Differences in cell proportions between the subiculum and CA1 in cases where cells are classified by the number of θ-phase–based fields: ‘SF’ for single field, ‘MF’ for multiple fields. **(C)** Comparison of mean firing rates of individual θ-phase–based fields for SF and MF cell types between the subiculum and CA1. **(D)** Comparison of field widths of individual θ-phase–based fields for SF and MF cell types between regions. \*\*\**p* < 0.001. st box, start box; fw, food well; SUB, subiculum.

Among cells exhibiting at least one θ-phase–based field (n = 135 in the subiculum; n = 168 in CA1), single place fields (SF cells) were more prevalent in CA1 (74.4%) compared to the subiculum (31.1%), whereas multiple place fields (MF cells) were more common in the subiculum (68.9%) than in CA1 (25.6%) (χ^2^(_2_) = 196.84, *p* < 0.001; chi-squared test) (**Fig. 2B**). Notably, a significant proportion of subicular MF cells had more than two subfields, whereas most CA1 MF cells typically exhibited only two subfields. The average firing rates of individual subfields were higher in SF cells than in MF cells in both regions (**Fig. 2C**). A two-way ANOVA with the number of fields and region as factors revealed a significant effect of the number of fields on average firing rates (F(_1,493_) = 37.45, *p* < 0.001) but no effect of region (F(_1,493_) = 2.31, *p* = 0.13).

A post hoc comparison showed that the average firing rates of individual fields of SF cells were significantly higher than those of MF cells in both regions (t(_272_) = 3.69, *p* < 0.001 in the subiculum; t(_218_) = 5.42, *p* < 0.001 in CA1). A comparison of fields between the subiculum and CA1 showed no significant difference in average firing rates (t(_165_) = 0.75, *p* = 0.454 for SF cells; t(_325_) = 1.573, *p* = 0.117 for MF cells). The field width differed significantly between SF and MF cell types (F(_1,493_) = 35.65, *p* < 0.001), a difference that was more prominent in the subiculum (t(_272_) = 5.63, *p* < 0.001) than in CA1 (t(_218_) = 1.96, *p* = 0.051) (**Fig. 2D**). As previously reported, place fields of subicular cells were larger than those of CA1 cells (F(_1,493_) = 35.65, *p* < 0.001), even when both displayed the same number of fields (t(_165_) = 6.99, p < 0.001 for SF cells; t(_325_) = 3.72, p < 0.001 for MF cells; Bonferroni-corrected two-sample t-test).

The θ-phase–based field identification method enabled us to identify task-demand–associated (TDA) rate modulation in the subiculum and CA1 (*34*) (**Fig. 2A**). In the VSM task, visual scenes and upcoming choice arms were defined as TDA variables. We quantified the strength of rate modulation for each θ-phase–based place field using the scene selectivity index (SSI) for visual scene pairs associated with the same rewarding arm: that is, SSI_L_ for the zebra-bamboo pair and SSI_R_ for the pebbles-mountain pair. Additionally, we calculated a choice selectivity index (CSI) to measure rate modulation strength based on the upcoming arm choice (see Material and Methods for details). A positive selectivity index indicated selective firing for the zebra scene, pebbles scene or left-arm choice, whereas a negative index indicated selective firing for the bamboo scene, mountain scene, or right-arm choice. A comparison of the distribution of selectivity indices of all θ-phase–based place fields showed that the levels of selectivity were similar between the subiculum and CA1 for all three task demands (*p* = 0.633 for SSI_L_; *p* = 0.199 for SSI_R_; *p* = 0.642 for CSI; Kolmogorov-Smirnov test; **Fig. S3A**).

We assessed the extent of rate modulation by comparing the absolute values of selectivity indices that measured the scene- and choice-associated task demands (**Fig. S3B**). A two-way ANOVA showed significant differences between SF and MF cells for all three types of task demands (F(_1, 299_) = 5.84, *p* < 0.001 for SSI_L_; F(_1, 298_) = 4.53, *p* < 0.001 for SSI_R_, F(_1,292_) = 1.98, *p* = 0.049 for CSI), but no significant differences between subiculum and CA1 (F(_1, 299_) = 1, *p* = 0.32 for SSI_L_; F(_1, 298_) = 0.15, *p* = 0.88 for SSI_R_, F(_1,292_) = 0.53, *p* = 0.6 for CSI). Specifically, SSI_L_ and SSI_R_ were larger in MF cells than in SF cells in both regions (SSI_L_: t(_133_) = -5.06, *p* < 0.001 in subiculum and t(_166_) = -3.3, *p* = 0.001 in CA1; SSI_R_: t(_133_) = -3.0, *p* = 0.003 in subiculum and t(_165_) = -3.42, *p* < 0.001 in CA1). However, CSI was not significantly higher in MF cells than SF cells in either the subiculum (t(_132_) = -1.81, *p* = 0.073) or CA1 (t(_160_) = -0.9, *p* = 0.369; Bonferroni-corrected two-sample t-test).

In summary, the overall neural firing characteristics were consistent between the subiculum and CA1 despite some differences between SF and MF cells. Importantly, our results demonstrate that cells in the subiculum and CA1 represent TDA variables (i.e., visual scenes and upcoming choice arms) in the VSM task through rate modulation at the level of θ-phase–based fields.

### Enhanced Reactivation of TDA Cells During Awake SWRs in the Subiculum Compared with CA1

Upon validating that place cells in the subiculum and CA1 exhibit rate modulation in response to task demands, we investigated whether the ensemble of cells simultaneously reactivated within an awake SWR event showed bias toward specific task conditions, such as a particular visual scene stimulus or choice arm. If most cells in a SWR-reactivated ensemble exhibited higher activity for a common task variable, we considered the ensemble to be reactivated selectively for that task variable during the SWR event. We termed the reactivation of cells exhibiting significant correlations with task demands, ‘TDA reactivation.’

We defined TDA reactivation using the following criteria. First, the cell ensemble should be present within the SWR boundaries. Then, we parsed the overall firing rate maps of the cells into individual θ-phase–based subfields to measure the selectivity indices of those fields (**Fig. 3A-C**). Fields with low selectivity indices (between -0.1 and 0.1) were excluded from the analysis. Cells with low selectivity indices for all subfields (1 subicular cell and 5 CA1 cells) were further excluded from the analysis. Subsequently, these field-based firing rate maps were reconstructed for various task demands (i.e., zebra vs. bamboo scene, pebbles vs. mountain scene, left vs. right choice arm) (**Fig. 3D-F**).

**Fig. 3.**
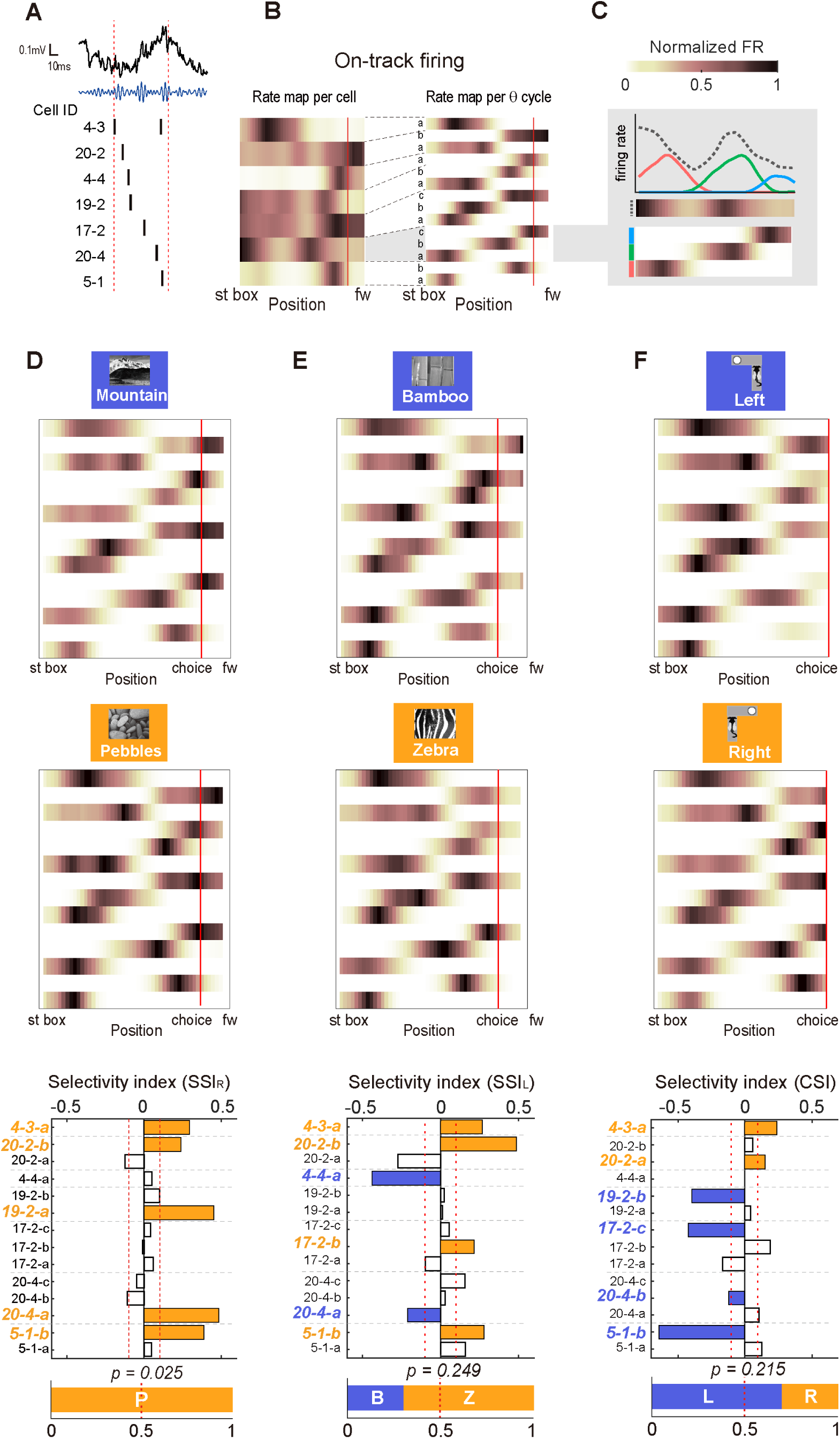
TDA bias in neuronal ensemble reactivation during SWRs. **(A)** An example of a subicular SWR during an ITI. Leftmost column: Raw LFP trace of a SWR event (solid black line; time and LFP amplitude indicated by the scale bar at the upper left) and 150-250 Hz band-pass filtered LFP trace (solid blue line). Dotted vertical red lines indicate the SWR boundaries. Raster plot under the traces shows spiking activity of the cell ensemble coactivated during the SWR. Cells are sorted based on the order of their first spike time. The numbers on the left side of the raster plot indicate the cell ID. **(B)** Cell-based–firing rate map (*rate map per cell*, left) and θ-phase–based rate map (*rate map per θ cycle*, right) constructed using all trials. Firing rates in each map are normalized to the cell’s maximum firing rate. Red solid line denotes choice point. **(C)** On-track firing activity of an example cell (18–3), presented as a line plot and a heatmap (normalized to peak firing rate). Gray dotted line, overall activity; solid colored lines, multiple θ-phase–based fields. **(D)** *Top and Middle*: θ-phase– based rate maps reconstructed according to right-arm–choice trials associated with the mountain versus pebbles scene. The rate map of each subfield is normalized to the maximum rate for that particular scene or choice condition. *Bottom*: Distribution of SSI_R_ for individual subfields. Each bar corresponds to a subfield in the rate map shown in the left panel. Dashed gray lines divide subfields from different cells. Empty bars represent selectivity indices that do not meet the criterion (dashed red lines; between -0.1 and 0.1) or that are not the highest within the cell. The bar graph at the bottom shows the proportion of cells in the reactivated ensemble that are selective for certain scenes or choice conditions. St box, start box; choice, choice point; fw, food well of the track; Z, zebra scene; B, bamboo scene; P, pebbles scene; M, mountain scene; L, left choice arm; R, right choice arm. **(E)** *Top and Middle*: θ-phase–based rate maps reconstructed according to left-arm–choice trials associated with the bamboo versus zebra scene. *Bottom*: Distribution of SSI_L_ for individual subfields. **(F)** *Top and Middle*: θ-phase–based rate maps reconstructed according to trials associated with left versus right choices. *Bottom*: Distribution of CSI for individual subfields.

To quantify TDA rate modulation at the ensemble level, we calculated a cell’s representative selectivity index for each task demand by finding the maximum value of the absolute selectivity indices among all task-demand–selective subfields of the cell (yellow or blue bars in **Fig. 3D-F**). For example, the subicular SWR event illustrated in **Fig. 3D-F** includes the reactivation of an ensemble whose scene-selective fields exhibited higher firing rates for the pebbles scene compared with the mountain scene. The bias toward the pebbles scene was also verified by the distribution of selectivity indices, with all cells’ representative selective indices showing positive values (yellow bars, *p* = 0.025) (**Fig. 3D**). On the other hand, at the same SWR, the distribution of selectivity indices for the Left scene pair had 67% of positive values (yellow bars), and a binomial test for this showed that there was no significant bias towards the zebra scene (*p* = 0.249) (**Fig. 3E**). In addition, the distribution of selectivity indices for the choice pair was 67% negative (blue bars), and a binomial test for this also showed no significant bias towards the left choice (*p* = 0.215) (**Fig. 3F**).

Our results revealed that a significant proportion of SWRs in both regions contained cell ensembles demonstrating substantial TDA reactivations for specific scenes or arm choices (**Fig. 4A** and **Fig. S4**). Notably, a significantly larger proportion of SWRs in the subiculum (61.5%) exhibited TDA reactivation compared to those in CA1 (41.5%) (χ^2^(_1_) = 170.9, *p* < 0.001; chi-squared test) (**Fig. 4B**). Additionally, p-values from binomial tests were significantly lower in the subiculum than in CA1 across all three task demands (Z = -12.57, *p* < 0.001 for SSI_L_: Z = -5.81, *p* < 0.001 for SSI_R_; Z = -7.48, *p* < 0.001 for CSI; Wilcoxon rank-sum test) (**Fig. 4C**), indicating a higher tendency for co-reactivation of cells encoding identical task demands during SWRs in the subiculum than in CA1.

**Fig. 4.**
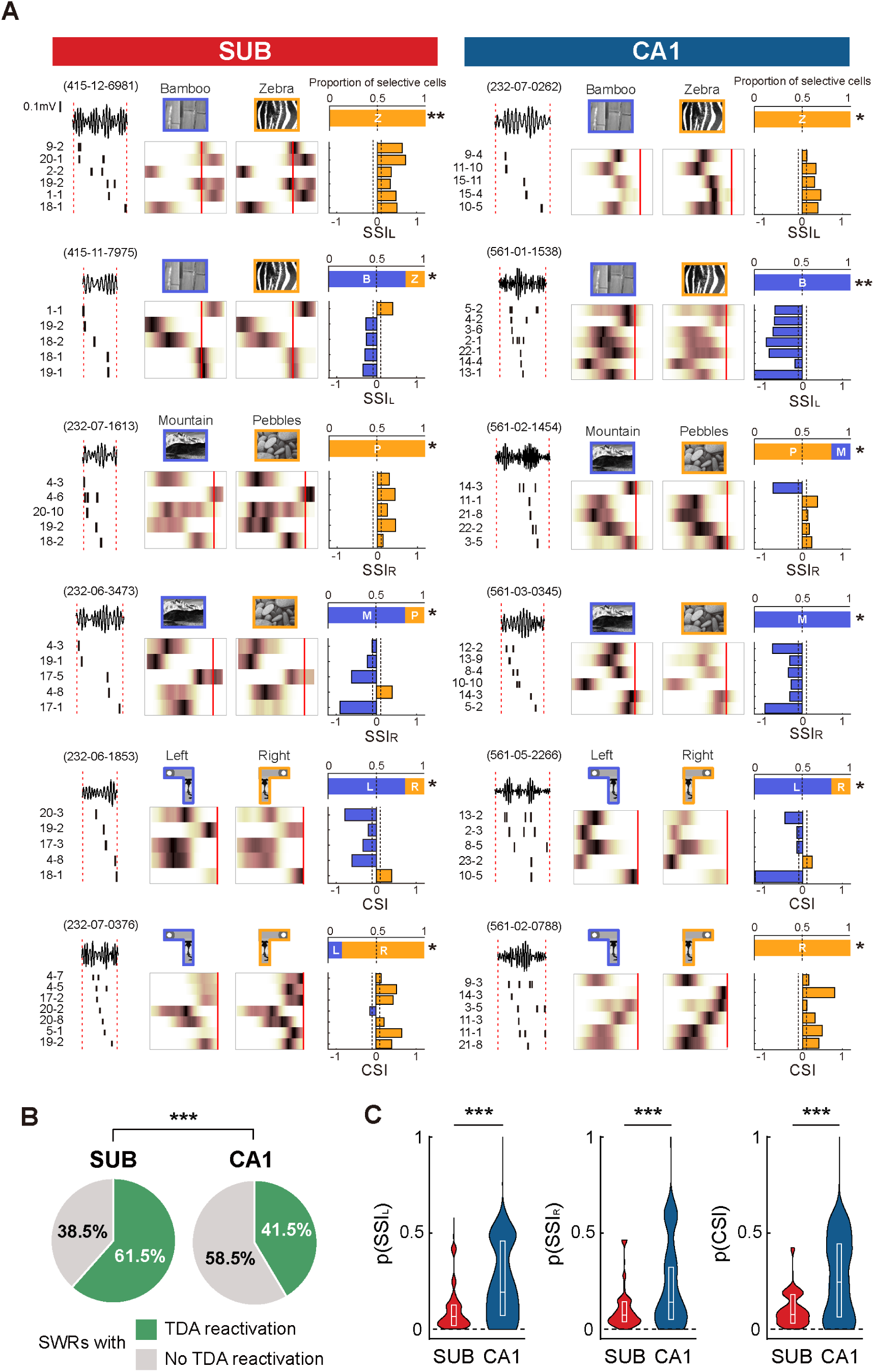
TDA reactivation in SWRs is more prevalent in the subiculum than in CA1. **(A)** Example SWRs with TDA reactivations of the subiculum (left) and CA1 (right). In each example, the leftmost part shows a filtered LFP trace and spiking activity of a cell ensemble within SWR boundaries. The middle colormaps represent the on-track firing rate map of θ-phase–based subfields from which representative selectivity indices were selected for each task condition. For the example SWRs with choice reactivation, ‘arm’ sections of T-maze, from choice point to food well, are excluded from display since these sections are not used to calculate choice selectivity index. The top right graph depicts the proportion of cells in the reactivated ensemble that are selective for certain trial conditions. The statistical significance of ensemble-level selectivity bias, calculated using a binomial test, is indicated as asterisks. Bottom right bar graphs are representative selectivity indices for individual cells, color-coded by trial condition. **(B)** Proportion of SWRs with significant TDA reactivation in each region. **(C)** Distributions of *p*-values obtained with the binomial test. In the inset box plot, the center line indicates median value, and top and bottom lines denote 1^st^ and 3^rd^ quartiles, respectively. \**p* < 0.05, \*\**p* < 0.01, \*\*\**p* < 0.001. St box, start box; choice, choice point; fw, food well; Z, zebra scene; B, bamboo scene; P, pebbles scene; M, mountain scene; L, left choice arm; R, right choice arm; SUB, subiculum.

TDA reactivation also occurred while the rats slept before and after the recording session (**Fig. S5A**). During the pre-sleep period, a significantly larger proportion of subicular SWRs (46%) exhibited TDA reactivation compared to SWRs in CA1 (17.8%) (χ^2^(_1_) = 98.1, *p* < 0.001; chi-squared test) (**Fig. S5B**). Similarly, in the post-sleep period, a significantly larger proportion of subicular SWRs (53.3%) showed TDA reactivation compared to SWRs in CA1 (32.7%) (χ^2^(_1_) = 106, *p* < 0.001; chi-squared test) (**Fig. S5C**).

These findings demonstrate that cell assemblies representing crucial TDA variables, such as visual scenes and upcoming arm choices, are reactivated during SWR events. More importantly, TDA neural reactivation is more prevalent and robust in the subiculum than in CA1, during both sleep and awake periods.

### Simultaneous Representation of Diverse Task Demands During TDA Neural Reactivation

Cells in the subiculum have demonstrated their superior capability of representing multiple TDA variables, such as scene and arm choice, in the VSM task, compared with those in the CA1 region (*34*). We further investigated whether different task demands could be simultaneously represented by cell ensembles during SWR events. **Fig. 5A** exemplifies SWR events wherein a scene or choice variable (or both) was represented during neural reactivations. Most SWRs exhibiting significant TDA reactivations showed scene selectivity only (68.4% in the subiculum; 65.6% in CA1), whereas some SWRs demonstrated choice-arm–selective reactivations exclusively in both regions (13% in the subiculum; 19.6% in CA1) (**Fig. 5B**). Notably, a subset of SWRs reactivated cell ensembles that represented both visual scene and choice-arm variables simultaneously, and these SWRs were more prevalent in the subiculum (18.5%) compared to CA1 (14.7%) (χ^2^(_1_) = 5.8, *p* = 0.016; chi-squared test).

**Fig. 5.**
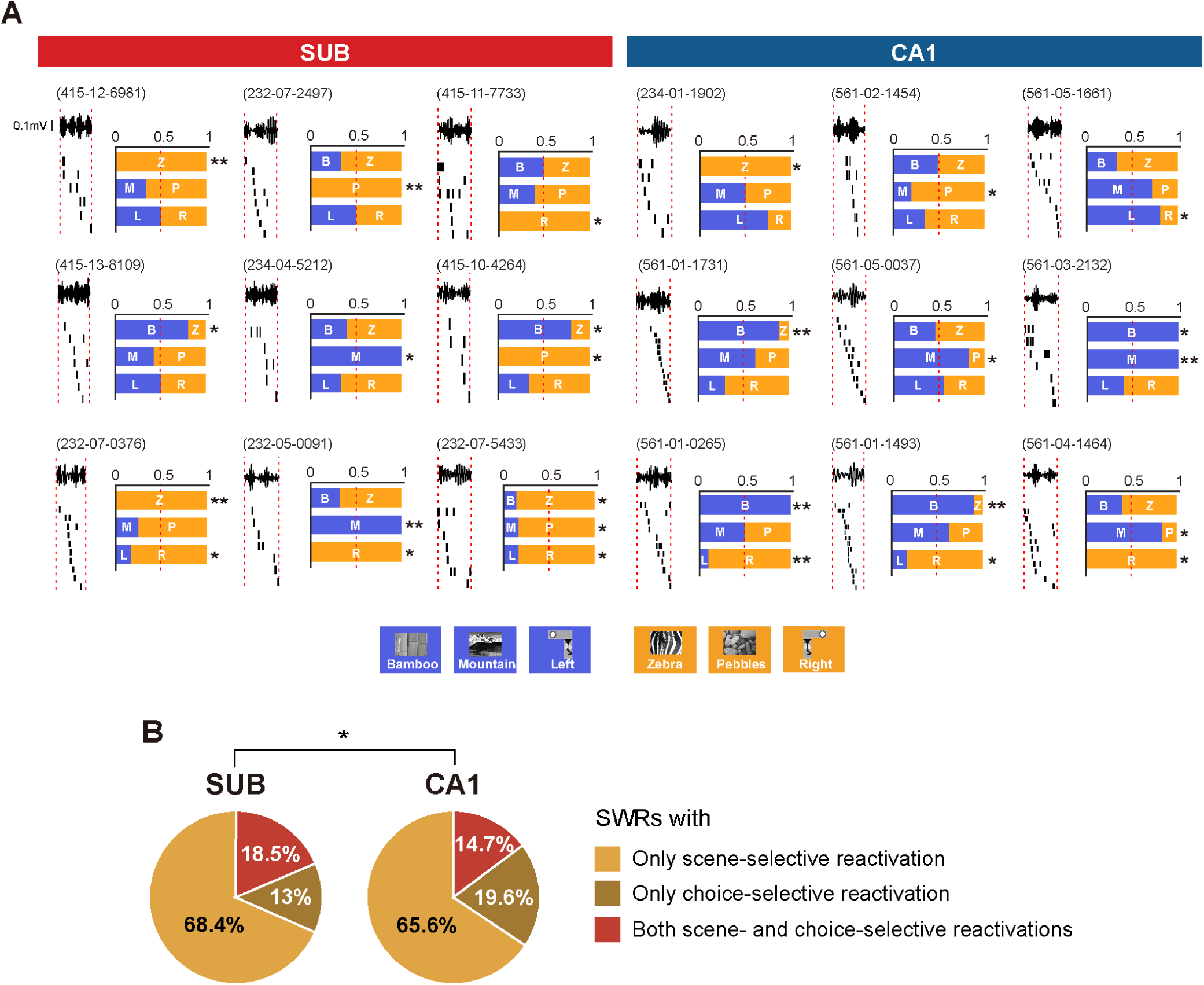
SWRs representing different types of TDA information are more prevalent in the subiculum than in CA1. **(A)** Example SWRs with scene-, choice- or both scene- and choice-selective reactivation in the subiculum (left panel) and CA1 (right panel). In each example, the leftmost graphic displays a filtered LFP trace and spiking activity within the SWR boundaries. Horizontal bar plots on the right indicate the proportions of cells in the reactivated ensemble that are selective for each scene or choice information. Asterisks indicate the significance of the proportional difference obtained from binomial tests. **(B)** Proportion of SWRs that exhibited scene-, choice- or both scene- and choice-selective reactivation within each region. \**p* < 0.05, \*\**p* < 0.01, \*\*\**p* < 0.001; Z, zebra scene; B, bamboo scene; P, pebbles scene; M, mountain scene; L, left choice arm; R, right choice arm; SUB, subiculum.

Our findings indicate that neural ensembles reactivated during SWR events do not simply represent a single type of task demand; rather, they often encode multiple types of TDA variables (i.e., visual scenes and upcoming choice arms) within the same SWR event. Furthermore, reactivations encompassing heterogeneous task demands occurred more frequently in SWRs recorded from the subiculum than from CA1. These results support our previous findings that subicular MF cells represent diverse task demands through their multiple θ-phase–based subfields (*34*).

### θ-Phase–Based MF Cells Drive TDA Reactivation in the Subiculum

Considering the superior capability of subicular MF cells to process heterogeneous TDA variables, such as visual scene and choice-arm information, compared with CA1 MF cells (*34*), we theorized that the increased frequency and diversity of TDA reactivations in the subiculum might be linked to the higher proportion of MF cells in this area relative to CA1.

We found that heightened neural activity in certain subicular MF cells coincided with TDA reactivations during SWR events, whereas the activity of subicular SF cells remained relatively stable, irrespective of the presence of TDA reactivations during SWRs. In contrast, SF cells in CA1 showed an increase in firing rates during SWRs while exhibiting TDA reactivations, unlike their MF counterparts (**Fig. 6A**). In support of this observation, the proportion of subicular MF cells within a reactivated ensemble during SWR events was larger, particularly when TDA reactivations were present (Z = -15.05, *p* < 0.001) (**Fig. 6B**). However, there was no similar trend in CA1 (Z = 0.26, *p* = 0.8; Wilcoxon rank-sum test).

**Fig. 6.**
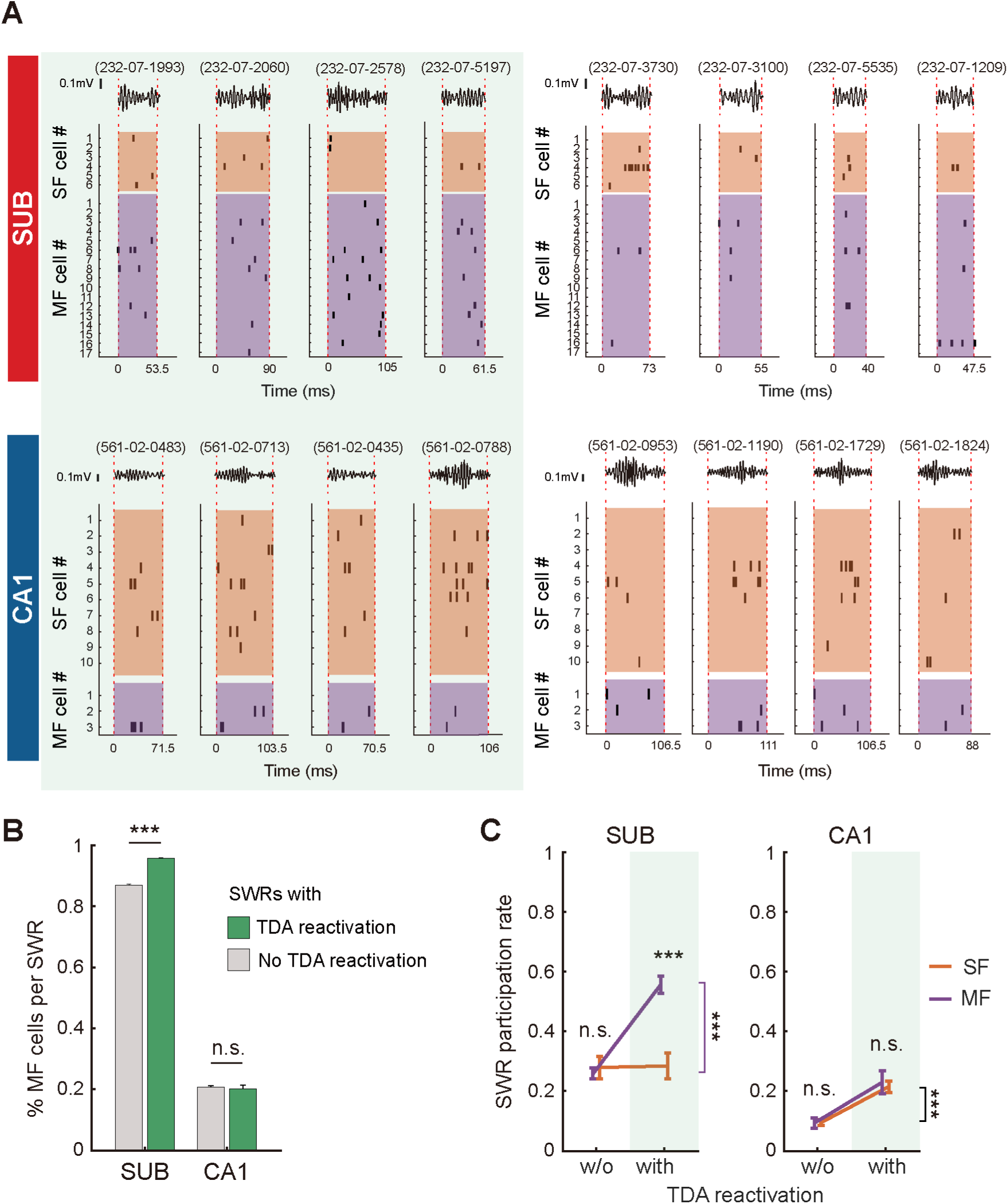
Contribution of subicular MF cells to TDA reactivation. **(A)** Example SWRs in the subiculum and CA1, with (left shaded area) and without (right) significant TDA reactivation. Filtered LFP traces are displayed, and the spiking activities of the reactivated cell ensembles are sorted by cell type (i.e., SF and MF cells). **(B)** Proportion of MF cells per SWR event. Data are presented as means ± SEM. **(C)** Comparison of SWR participation rates for individual cells between cell types and SWR types (i.e., with or without TDA reactivation) within each region. \*\*\**p* < 0.001; SUB, subiculum. w/o, SWRs without TDA reactivation.

To mitigate the potential influence of varying MF cell numbers between the regions, we analyzed the participation rates of each cell type in SWR events. Engagement of subicular MF cells in SWRs was higher in cells with TDA reactivations than in those without TDA reactivations (t(_176_) = 9.12, *p* < 0.001). In contrast, subicular SF cells did not exhibit such difference in behavior (t(_72_) = 0.08, *p* = 0.93) (**Fig. 6C**). Both MF and SF cell types in CA1 demonstrated a marked increase in participation rates in SWRs with TDA reactivations (t(_81_) = 3.28, *p* = 0.002 for MF cells; t(_247_) = 5.68, *p* < 0.001 for SF cells). A comparison of participation rates in SWRs with TDA reactivations showed that subicular MF cells exhibited significantly higher rates compared to SF cells (t(_115_) = -4.88, *p* < 0.001), whereas SWR participation rates were not significantly different between MF and SF cells in CA1 (t(_162_) = 0.36, *p* = 0.72). For TDA reactivations in the absence of SWRs, differences in participation rate between MF and SF cells were insignificant in both the subiculum and CA1 (t(_133_) = -0.66, *p* = 0.51 in the subiculum; t(_166_) = 0.09, *p* = 0.93 in CA1; Bonferroni-corrected two-sample t-test). Moreover, SWR-based TDA reactivations significantly decreased when the analysis was conducted using place fields defined by firing rates, particularly in the subiculum, where most cells exhibited broad single fields (χ^2^(_1_) = 23.7, *p* < 0.001; chi-squared test for proportions of TDA reactivation in the subiculum and CA1) (**Fig. S6**).

Our results indicate that subicular cells with multiple θ-phase–based subfields may be pivotal in driving the frequent and robust reactivation of TDA variables during SWRs.

### Functional Separation of Neural Ensemble State during SWRs in the Subiculum, but not in CA1

We further investigated latent variables that could potentially separate the neural ensemble states during the SWR-based reactivations (with or without TDA reactivation) to differentiate the subiculum further from CA1. For this purpose, we used the t-distributed stochastic neighbor embedding (t-SNE) algorithm to reduce the collective reactivation pattern of many neurons to three dimensions. This nonlinear dimensionality reduction method extracts the patterns embedded in the high-dimensional structure of ensemble reactivation into a low-dimensional structure. This will generate a low-dimensional manifold from our data in which SWRs sharing a common reactivated ensemble appear clustered together. In this analysis, we focused on measuring the TDA information represented in the reactivated neural ensemble across all SWRs, rather than just TDA reactivation.

Our analysis revealed the differences between the subiculum and CA1 in the neural manifold of the reactivated ensembles during SWR (**Fig. 7**; see Materials and Methods for details). Specifically, the neural manifold of subicular SWRs showed two clusters that were relatively well-separated, whereas the neural manifold of CA1 SWRs did not. This difference was further emphasized when colored by the ratio of reactivated cell types within the reactivated ensemble. In the Subicular SWR (**Fig. 7**, 1^st^ and 2^nd^ columns), the reactivated ensembles with a slightly higher proportion of Pebbles scene-selective cells (colored in light blue) and the reactivated ensembles with a slightly higher proportion of Mountain scene-selective cells (colored in light green) were located far away from each other, in two distinct clusters, whereas in the CA1 SWRs (**Fig. 7**, 3^rd^ and 4^th^ columns) they were relatively close to each other, seemingly forming a single cluster with a continuous color gradient. This suggests that distinct neural ensembles were co-reactivated in the two SWRs of the subiculum, while neural ensembles that highly overlapped with each other were co-reactivated in the two SWRs of CA1.

**Fig. 7.**
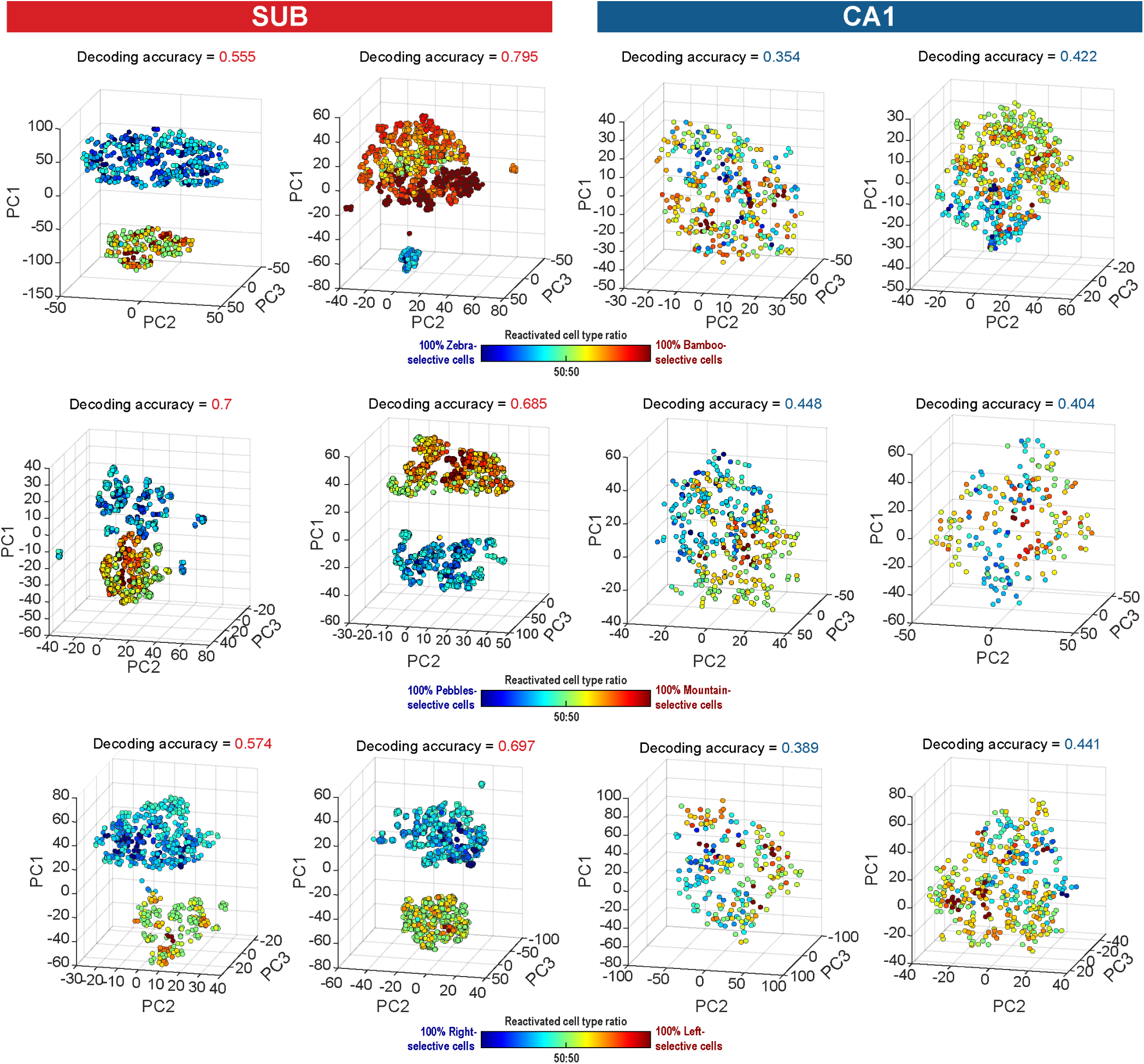
Neural ensemble state is separated in subicular SWRs along the selectivity distribution of their reactivated cells, compared to the continuous pattern of CA1 SWRs. The neuronal manifold during sharp-wave ripple (SWR) events is illustrated in three-dimensional space for the subiculum (1^st^ and 2^nd^ columns) and CA1 (3^rd^ and 4^th^ columns). Each individual data point represents the activity of a neuronal ensemble during a specific SWR. Data points are color-coded based on the ratio of selective cell types reactivated during each SWR (e.g., scene-selective cells for “Zebra” versus “Pebbles” for the 1^st^ row; scene-selective cells for “Pebbles” versus “Mountain” for the 2^nd^ row; choice-selective cells for “Left” versus “Right” for the 3^rd^ row). SUB, subiculum

To quantify the degree to which the clustering of neural manifolds can be explained by a label based on the reactivated cell type ratio in the reactivated ensemble, we trained a K-NN clustering model using 20% of the data from each manifold and evaluated the decoding accuracy of the model. The decoding accuracy of the k-nearest neighbors (K-NN) clustering model with the output label of reactivated cell type ratio was significantly higher for the neural manifold of subicular SWR (mean = 70.51%, median = 71.22%) than for that of CA1 SWR (mean = 42.33%, median = 43.15%) (Z = 4.78, *p* < 0.001; Wilcoxon rank-sum test).

Taken together, these results show that the reactivated neural ensemble state during subicular SWR converges to two distinct clusters according to the ratio of reactivated cell types. This implies that even for subicular SWRs without TDA reactivations, TDA information strongly predicts which neuronal ensembles are co-reactivated in the SWR.

### More Prevalent Spatial Reactivations during SWRs in CA1 than in the Subiculum

We next investigated whether spatial reactivation, that is, spatial replay of place fields (*6–8*), is also found during awake SWR events in both regions. To analyze spatial reactivation, we utilized the complete on-track firing patterns of cells instead of θ-phase–based fields owing to the challenge of assigning spikes during SWR events to specific fields. The posterior probability of locations on the maze was calculated by implementing Bayesian decoding, generating a linear regression strength value (R^2^) (**Fig. S7A-C**; see Material and Methods for detailed methodology).

Our observations revealed similar patterns of spatial reactivation in both the subiculum and CA1 regions (**Fig. S7D**). Interestingly, the percentage of SWRs exhibiting spatial reactivation was significantly higher in CA1 (20.0%) than in the subiculum (6.0%) (χ^2^(_1_) = 729.04, *p* < 0.001; chi-squared test) (**Fig. S7E**). Moreover, the linear regression strength was notably greater in CA1 than in the subiculum (Z = -19.81, *p* < 0.001; Wilcoxon rank-sum test) (**Fig. S7F**). Notably, only a small fraction of SWRs in CA1 (3.4%) and the subiculum (1.2%) demonstrated both spatial and TDA reactivations, suggesting that these two types of reactivations occur separately in most instances.

Furthermore, the presence of spatial reactivations led to a decrease in the proportion of MF cells within the reactivated ensemble in the subiculum, but not in CA1 (t(_5897_) = 21.47, *p* < 0.001 in the subiculum; t(_1889_) = 0.54, *p* = 0.59 in CA1; two-sample t-test) (**Fig. S7G**). This decline in the proportion of MF cells during spatial reactivations supports the independent nature of spatial and TDA reactivations, aligning with prior studies highlighting the pivotal role of subicular MF cells in TDA reactivations.

These findings indicate that, while spatial reactivation, often referred to as ‘replay’, occurs in subicular SWRs, its frequency and robustness in this region are lower compared with CA1.

## Discussion

In the current study, we found that the reactivation of TDA variables during SWRs was more pronounced in the subiculum than in CA1. Additionally, the incidence of SWRs representing diverse TDA variables was higher in the subiculum than in CA1. Specifically, subicular neurons with multiple θ-phase–based fields demonstrated increased involvement in TDA reactivation. In addition, the neural state of the reactivated ensembles in the subicular SWR showed more clustered manifold, separated by task-related variables, implying that TDA information of individual neurons strongly influences neuronal co-reactivation during SWRs in the subiculum. On the other hand, while spatial replays were also identified in the subiculum, they occurred at a lower frequency than in the hippocampus. Crucially, these spatial replays in the subiculum were observed independently of TDA reactivations. Collectively, our results highlight the pivotal role of the subiculum in facilitating the transmission of crucial task-relevant information between the hippocampus and cortical regions during hippocampal-dependent memory tasks.

### Multiple θ-Phase–Based Place Fields Drive Frequent and Diverse TDA Reactivations in the Subiculum

Literature has reported a correlation between neural activity in the subiculum and hippocampal SWRs (*28,29*). Bohm et al. (2015) found that subicular neurons exhibited varying responses to hippocampal SWRs based on their intrinsic firing patterns. Utilizing whole-cell current-clamp recordings in both awake, head-fixed mice and in vitro settings, they classified subicular pyramidal cells as either bursting or regular spiking, reporting that firing rates of bursting cells increased around the peak of the ripple, whereas those of regular spiking cells decreased. Similarly, Kitanishi et al. (2021) observed ripple-associated firing rate modulation in subicular excitatory neurons, noting that the modulation varied depending on the efferent regions to which the neurons projected. Specifically, neurons projecting to the anteroventral thalamus were suppressed following hippocampal SWRs, whereas those projecting to the nucleus accumbens (NAc) or medial mammillary body were activated. These findings indicate that subicular neurons are responsive to hippocampal SWRs, although the specific type of information represented during these events in the subiculum has remained unclear.

In this study, we examined the reactivation of TDA information in the subiculum during SWRs to determine the type and extent of information represented by subicular neurons compared with hippocampal cells in a VSM task. Notably, we developed a novel method for defining subicular place fields based on spiking phases relative to local theta oscillations. We found that two-thirds of the recorded cells in the subiculum were classified as having multiple θ-phase–based place fields (MF), displaying firing properties similar to those of typical place fields in CA1. These MF cells exhibited greater selectivity for task variables than SF cells in the subiculum and encoded heterogeneous TDA variables through their multiple subfields.

We hypothesized that subicular MF cells play a critical role in hippocampal-dependent associative memory tasks. In support of our hypothesis, the unique firing characteristics of subicular MF cells significantly influenced TDA reactivations during SWRs. Specifically, the participation rate of subicular MF cells in SWRs was selectively increased when TDA information was reactivated (**Fig. 6C**), leading to the reactivation of heterogeneous task-related information within a single SWR event (**Fig. 5**). These results not only confirm our previous findings but also suggest a functional diversity of subicular principal neurons in episodic memory.

We further observed that the participation rates of SF cells and MF cells in TDA reactivations varied only in the subiculum, even though the overall firing profiles of MF cells on the track were similar between the subiculum and CA1 (**Fig. 2**). The selective involvement of subicular MF cells in TDA reactivations might be attributable to their orthogonal representations (*34*). This means that task-relevant variables showing the greatest selectivity differ across multiple fields of a single neuron. For example, one field might represent a visual scene, whereas another could represent an upcoming choice arm. In contrast, in most cases of CA1 MF cells, heterogeneous representations were driven by individual (single) fields. This suggests that SF cells and MF cells in CA1 do not functionally differ in representing TDA variables, which explains why the participation rates in TDA reactivation were similar between the two cell types in the CA1.

To examine TDA reactivations, we chose the subfield with the highest selectivity index among multiple fields as the representative field for each TDA variable. A limitation of this analysis is that it is difficult to assume that a spike generated during a SWR originates from a specific field. The current analysis assumed that the spike signals the TDA variable with the highest selectivity index, which is most strongly represented by the cell. This issue also arises in spatial reactivation analyses of place cells with multiple place fields. Particularly in large spaces or environments with many repetitive compartments, a significant portion of place cells exhibit multiple place fields in the hippocampus (*37–40*). Future studies are needed to develop a method to determine which specific field or information a spike represents during SWR signals to enable a more accurate investigation of the reactivated content.

### TDA Reactivations During SWRs Operate Independently of Spatial Replays

Numerous studies have consistently shown that hippocampal place cells are sequentially reactivated within a brief time window during a SWR event, reflecting previously experienced trajectories (*6–8*). Additionally, recent research indicates that the quantity of reward or value associated with different locations modulates spatial reactivations both quantitatively and qualitatively in the hippocampus (*10–14*). For instance, replayed trajectories tend to favor locations associated with greater rewards (*11*) or higher value rewards (*14*). Similarly, the frequency of SWRs and spatial replays adjusts to reflect the relative magnitude of reward between two ends of a linear track (*10*). Furthermore, studies have demonstrated that differences in firing rate related to an animal’s running direction or environmental context are preserved in spatial replay patterns within hippocampal SWRs (*18,19*).

While the aforementioned studies focused on how task-relevant variables influence spatial reactivations during SWRs, the present study highlights the neural reactivation of TDA information during SWRs, which occurs independently of spatial reactivations. The likelihood of a neuron representing both spatial and task-relevant factors simultaneously is very low in both the hippocampus and subiculum. Specifically, we found that, whereas approximately 40% of SWRs in CA1 involved TDA reactivations and 20% involved spatial reactivations, only 3.4% exhibited both spatial and TDA reactivations (**Fig. 4B** and **S7E**). This phenomenon was more pronounced in the subiculum, where just 1.2% of SWRs showed concurrent spatial and TDA reactivations. Moreover, in the subiculum, the proportion of MF cells per SWR increased during TDA reactivations (**Fig. 6B**) and decreased during spatial reactivations (**Fig. S7G**).

Our findings suggest that TDA variables, such as surrounding visual scenes and upcoming choice arms, are reinstated during SWRs in both the hippocampus and subiculum through the recruitment of neural ensembles with similar selective firing patterns, independent of their spatial representations on the track. This discovery opens new avenues for investigating SWR-related reactivation patterns of neurons recorded from various regions of the hippocampal formation (e.g., subiculum, intermediate hippocampus), whose spatial correlates are not as precisely tuned as the typical place cells found in the dorsal hippocampus.

### The Functional Significance of TDA Reactivations in the Subiculum

Previous research has shown that hippocampal SWRs are transmitted to the granular retrosplenial cortex via the subiculum (*26*). However, we found that fewer TDA reactivations occurred in CA1 than in the subiculum (**Fig. 4B**), although the proportion in CA1 was still significant. Building on this evidence, we suggest that the subiculum not only helps hippocampal SWRs propagate to cortical regions during hippocampal-dependent memory tasks, but it also amplifies and transmits signals carrying TDA variables to cortical areas. Another possibility is that TDA reactivations could originate within the subiculum itself and then propagate to the hippocampal formation, as SWRs independently generated in the subiculum have been reported (*27*). However, since back-projection from the subiculum to CA1 is known to function under limited task demands, such as object-in-place memory (*41,42*), this hypothesis needs further testing.

Importantly, although TDA reactivations were observed during sleep in our task, TDA reactivations were observed more frequently in the awake state, specifically during behavioral tasks when rats were actively engaged. Unlike ripples during slow-wave sleep, which are thought to play a role in the consolidation of hippocampal episodic memory (*20,21,43,44*), awake ripples are considered important for active route planning or path simulation to reach goal locations in an environment (*45–47*). Disruptions of awake ripples have been shown to impair memory-guided navigational tasks (*21*). The TDA reactivations observed in the current study may also facilitate the prediction or simulation of visual scene stimuli as cues for upcoming choice arms or required responses to reach goal locations.

On the other hand, consistent with prior studies, spatial replay was predominant in CA1 (**Fig. S7**). Moreover, these spatial reactivations likely propagate while bypassing the subiculum. The poorer spatial tuning of subicular place cells compared with those in CA1 (*30–34*) may provide an intuitively appealing explanation for the relative rarity and diminished robustness of spatial replay in the subiculum. Some recent papers have argued that the subiculum and CA1 possess similar levels of spatial information (measured as information per second) (*29, 33*). Reports further indicate no significant difference in the decoding performance of an animal’s position between the two regions (*29*). Nevertheless, these analyses were based on neural activity recorded while the animal was running on a track, so whether the subiculum and CA1 have the same level of spatial representation during the short time window of a SWR remains unclear.

Overall, TDA reactivations are primarily driven by subicular MF cells that may receive convergent inputs from single place cells in CA1 and then transmit this information to cortical and subcortical regions associated with goal-directed behavior or motivation. This conclusion is supported by previous reports that TDA representations, such as the prospective coding of choice arm on a T-maze, are observed at particularly high rates in NAc-projecting neurons in the subiculum, and that the activity of these NAc-projecting neurons is enhanced during SWRs (*29*). Therefore, we propose that the subiculum and CA1 cooperate to organize neural network activity during SWRs by transmitting spatial or TDA variables in a complementary manner. Furthermore, the subiculum reactivates multiple types of TDA information within a single SWR event, potentially enhancing associative learning by integrating and multiplexing heterogeneous task-related factors, as suggested by our earlier study (*34*).

## Materials and Methods

### Subjects

The current study used five Long-Evans male rats, each weighing 350-400 g. Food was restricted to keep their body weight at 85% of their free-feeding weight and to maintain their motivation during the behavioral task, whereas water was available ad libitum. The rats were individually housed and kept on a 12-hour light/dark cycle. All procedures were carried out according to the guidelines of the Institutional Animal Care and Use Committee (IACUC) of Seoul National University (SNU-200504-3-1).

### Behavioral task

Our previous studies (*32,34*) provide a detailed description of experimental procedures, including the visual scene memory (VSM) task and the apparatus. Briefly, rats were required to make a left or right turn on the T-maze according to the patterned visual stimulus (i.e., visual scene) presented on the LCD monitors surrounding the arms. Before the initiation of each trial, the rat was placed in a start box enclosed by black walls (22.5 × 16 cm; height, 31.5 cm). Upon opening the start box’s guillotine door by the experimenter, the rat was exposed to one of four visual scene stimuli displayed on an array of three adjacent LCD monitors, signaling the onset of the trial. The rat then ran along the stem of the T-maze (73 × 8 cm) and had to choose the left or right arm (38 × 8 cm) at the end of the stem (i.e., ‘choice point’) based on the visual scene. If the rat chose the correct arm, it received a quarter piece of cereal reward (Kellogg’s Froot Loops) from the food well at the end of the arm. If it chose the wrong arm, no reward was given. After making a correct or incorrect choice, the rat was returned to the start box, and the door was closed to prevent the rat from seeing outside during the inter-trial interval (ITI) before the start of the subsequent trial. Four grayscale visual patterns – zebra stripes, bamboo, pebbles and mountain – were used for visual scenes. Of these four scenes, zebra stripes and bamboo patterns were associated with the left arm in all sessions, whereas pebbles and mountain patterns were associated with the right arm. These four visual scenes were presented in a pseudorandom sequence within a session. Data obtained from incorrect trials and subsequent ITIs were excluded from the analysis.

Before surgery, the rat learned two scene pairs sequentially (zebra vs. pebbles or bamboo vs. mountain), counterbalanced across rats, until it met the performance criterion for each pair (≥75% correct for each scene for two consecutive days, with 40 trials per session). Rats required an average of 2 weeks (13.4 ± 0.9 sessions, mean ± STE) to meet the performance criterion for both pairs.

### Hyperdrive implantation surgery and electrophysiological recording

Once the rat met the performance criterion, a hyperdrive equipped with 24 tetrodes and three additional reference electrodes was surgically implanted into the right hemisphere to cover an area from 3.2 to 6.6 mm posterior to bregma and 1 to 4 mm lateral to the midline. Following 1 week of recovery, the rat underwent retraining with two pairs of scene stimuli, in order, until it regained its presurgical performance levels, completing up to 160 trials. During this postsurgical retraining phase, the tetrodes were progressively lowered into the subiculum and CA1 at a rate of 40 to 160 μm per day. Once the optimal number of single units in the target regions was achieved, the main recording sessions commenced (123 ± 6 trials per session, mean ± STE).

Neural signals were transmitted through the headstage and tether attaching the electrode interface board of the hyperdrive to the data acquisition system (Digital Lynx SX; Neuralynx). Neural signals measured by tetrodes are amplified 1,000-10,000 times and digitized at 32 kHz using the Digital Lynx system, after which the signals were filtered at 600-6,000 Hz for spiking data and 0.1-1,000 Hz for LFP data.

### Histological verification of electrode tip positions

After all recordings were completed, an electrolytic lesion was performed on each tetrode (10 μA for 10 seconds) to mark the tip position. Twenty-four hours later, the rat was euthanized by inhalation of a lethal dose of carbon dioxide (CO_2_) and perfused transcardially. After post-fixation procedures, coronal brain sections (40 μm thick) were stained with thionin and using Timm’s method. Electrode tip positions were verified using a series of brain sections, considering the presurgical electrode configuration and electrolytic lesion marks (see our previous paper for details (*32*)). The anatomical boundaries of the subiculum and CA1 were established by reference to a rat brain atlas (*48*). Electrodes placed in the CA1-subiculum transition zone were excluded from further examination.

### Unit isolation and filtering

Single units were manually isolated using both commercial software (SpikeSort3D; Neuralynx) and a custom-written program (WinClust), based on the waveform parameters, peak amplitude and energy. Units qualified for analysis if they exhibited an average peak-to-valley amplitude greater than 75 μV and less than 1% of their spikes occurred within the refractory period (<1 ms). Fast-spiking units, characterized by an average firing rate greater than 10 Hz and an average waveform width less than 325 μs, were excluded from the analysis. Following these criteria, 192 and 282 complex-spiking units recorded in the subiculum and CA, respectively, were ultimately included in the current study.

### Construction of linearized firing rate maps

Position data collected during the outbound period of each trial (from leaving the start box to reaching the food well) were binned into 2-cm spatial bins. For each spatial bin between stem and arm, the significance of trace divergence (difference in animal position along the arm axis) between trials to the left arm and trials to the right arm was tested (two-sample *t*-test). A choice point was defined as a case where at least two consecutive spatial bins showed a significant trace difference. For the intersection (from the choice point to arms), the animal position was linearized to one-dimensional coordinates, considering both the animal’s x and y positions, and the spatial bins were fitted to a triangular shape along the animal’s trace. Then, a linearized firing rate map was created by dividing the number of spikes by the number of position data (recorded 30 times per second) for each bin. The linearized firing rate map was smoothed for display using an adaptive binning method (*49*).

### Identification of spiking theta phases

The recorded LFP data were down-sampled to 2 kHz and then filtered using a 3-300 Hz bandpass filter (third-order Butterworth filter using the *filtfilt* function in Matlab). For each tetrode, the power spectral density function was applied using the multi-taper method (Chronux ToolBox; Matlab), and the reference electrode with the strongest power in the high theta band (7-12 Hz) in each session and region (subiculum and CA1) was selected. A high theta band range was chosen to minimize artifacts (e.g., bumping noises). LFP data of the reference electrode were then filtered in the theta band (7-12 Hz). Spiking theta phases were obtained by parsing LFP data during the outbound running epoch on the track (speed > 20 cm/s). The theta phase of the reference tetrode at the moment when a spike occurred was adopted as the theta phase of the spike.

### θ-phase–based field identification

Place fields in the subiculum and CA1 were identified by applying the analytical method used in our previous study (*34*), which is based on the spiking phase relationship called theta phase precession (*50,51*). To identify a group of spikes consisting of a single cycle of theta phase precession from the entirety of spiking activity on the outbound journey on the T-maze, we used the density-based, non-parametric clustering algorithm, DBSCAN (*52*). First, the theta phases of spikes were plotted according to position on the outbound track for each complex-spiking cell. The algorithm then detected a cluster of spikes based on the predetermined parameters, distance (ε) between spike points and minimum number of points (N_min_) within that distance. In clustering, a core point was identified if there were more than N_min_ data points within a radius of ε. Points within this radius were termed border points. Points outside this radius without neighboring core points were classified as noise points. A cluster consisted of a core point and its border points, and intersecting clusters were regarded as a single entity. Clusters with insufficient spike count (<30) or irregular cluster shape were excluded, and the remaining clusters were used as ‘θ-phase–based place fields’ in rate modulation analysis. Only cells with at least one θ-phase– based field were used for SWR detection and reactivated ensemble analysis (CA1, n = 168 cells; subiculum, n = 135 cells). The width of each firing field was calculated by multiplying the number of bins within the place field by the spatial bin size (2 cm).

### Rate modulation analysis

To quantify differences in firing rate between trial conditions (i.e., visual scenes or upcoming choice arms) within a place field, we calculated the selectivity index (SI) according to the equation (1):

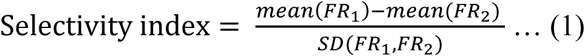

where FR_1_ and FR_2_ denote the in-field firing rates of the trials associated with two different conditions. For the selectivity index between choice directions (choice selectivity index [CSI]), FR_1_ was used for the in-field firing rate in left choice trials, whereas FR_2_ was used for right choice trials. CSI was computed based on spiking activity up to the choice point on the track, to isolate the influence of upcoming directions and minimize the impact of confounding variables such as varying positions. In addition, subfields were excluded from CSI calculation if their peak firing locations fell within track arms. For selectivity indices between visual scene stimuli, we computed two distinct scene selectivity indices (SSI): one for the pair of scenes associated with left arm choice (SSI_L_; for the zebra-bamboo scene pair) and another for the pair associated with right arm choice (SSI_R_; for the pebbles-mountain scene pair). These calculations included data from both the choice point and track arms. A field was excluded from further analysis for a certain condition if its selectivity index for that condition fell between -0.1 and 0.1. The representative selectivity index of a cell for each task condition was determined as the largest (negative or positive) value of the selectivity indices of all survived fields of the cell.

### Firing-rate–based field identification

The boundaries of a rate-based place field were identified as the first spatial bin at which the firing rate was 33% or less of the peak firing rate for two consecutive bins. If a local peak exceeding 50% of the maximum peak firing rate was found outside the previously defined firing field, it was considered the peak of another potential field. The boundaries of that field were then determined using the same method.

The spatial information score was computed according to the equation (2):

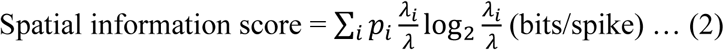

where *i* denotes each spatial bin, *p_i_* is the occupancy rate in the *i^th^* spatial bin, *λ_i_* is the mean firing rate in the *i^th^*spatial bin, and *λ* is the overall mean firing rate on the track (*49*). The peak and mean firing rates of cells were calculated from raw firing rate maps. Firing fields that met the following criteria were ultimately defined as ‘rate-based place fields’: (1) peak firing rate > 1 Hz, and (2) a spatial information score > 0.5.

### Sharp-wave ripple detection

SWRs were detected using LFP signals (filtered with a 150-250 Hz band-pass filter) recorded from representative tetrodes targeting the subiculum (for subicular SWRs) and CA1 (for CA1 SWRs). The tetrode exhibiting the highest ripple band power during the ITI was initially selected as the representative tetrode for each session and region. The Hilbert transform was then applied to obtain the envelope of the ripple band activity from this representative tetrode. Subsequently, SWR events were identified as segments exceeding 2 standard deviations (SDs) of the mean of the filtered signals. The boundaries of each SWR event were extended to include sections above the mean + 0.5 SD (computed for each session’s envelope during the ITI). In cases where two SWR events occurred within intervals shorter than 20 ms, the events were merged into a single event. SWRs with durations shorter than 40 ms or longer than 200 ms were excluded from the analysis. SWR events were further refined based on the number of cells reactivated during the event for TDA or spatial reactivation analysis. CA1 SWRs were characterized as events in which at least three CA1 cells were reactivated within the SWR duration. Similarly, subicular SWRs were identified as events where a minimum of three subicular cells were reactivated. The participation rate of each cell in SWRs was quantified as the ratio of SWRs that elicited at least one spike from the cell to the total number of SWRs recorded in the session.

### Identification of TDA reactivation

To determine the reactivation of TDA information by the cell ensemble during SWRs, we assessed whether the cells’ representative selectivity indices exhibited a significant bias toward a specific trial condition. To test this, we calculated the proportion of cells with a positive selectivity index and the proportion of cells with a negative selectivity index in the simultaneously recorded cell ensemble. We then conducted a binomial test to determine whether the proportion of positive/negative selective cells in the reactivated ensemble could occur by chance when compared to the ensemble-level proportion. Reactivated ensembles with *p*-values < 0.05 in the binomial test were defined as TDA reactivations.

### Identification of spatial reactivation

To examine spatial reactivation, we reconstructed the rat’s position on the track from the spiking activity in SWRs by applying a Bayesian decoding algorithm (*53,54*). In Bayesian decoding analysis, the duration of each SWR event was divided into 10-ms temporal bins. For individual bins with at least one spike, the conditional probability, P(x|n), that the animal is located at position x when the spike count is n is given by following formula (3):

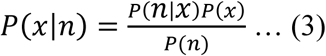

where P(n) is the probability that the spike count is n, and P(x) is the probability that the animal is positioned at x. If each cell, i, exhibited firing characteristics that follow a Poisson distribution and that all N cells were active independently, P(n|x) was calculated as (4):

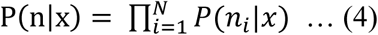

and P(x|n) was computed as (5):

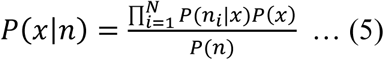

The strength of spatial reactivation was quantified as an R^2^ value, obtained through linear regression performed on the decoded spatial probability distribution across temporal bins. Statistical significance was also assessed by obtaining R^2^ values from shuffled data, created by randomizing the cell IDs of spikes fired during SWRs. The shuffling procedure was repeated 2,000 times, and the *p*-value was determined from the proportion of shuffled R^2^ values greater than the actual value. SWRs with p-values < 0.05 were considered to exhibit spatial reactivation.

### Dimension reduction and cluster quality analysis for reactivated ensemble state

We transformed each SWR into a binary vector with dimensions equal to the number of neurons, with each element of this vector representing the firing state of a particular neuron during the SWR (1: fired, 0: did not fire). These high-dimensional vectors were then reduced to three dimensions using the t-distributed Stochastic Neighbor Embedding (t-SNE) algorithm. The initial values for the t-SNE algorithm were obtained from a three-dimensional Principal Component Analysis (PCA), and subsequently modified according to the t-SNE algorithm to reflect the correlations between data points derived from the original dataset. The reduced three-dimensional data were then plotted in three-dimensional space, separated by session, so that each plot contained SWRs from only a single session.

Subsequently, we conducted clustering analysis of these 3D data points based on the distribution of neuronal ensemble types reactivated during the SWRs. For each data point, we employed the ratios of cell types reactivated during the SWR (e.g., zebra vs. bamboo scene-selective cells, pebble vs. mountain scene-selective cells, left vs. right choice-selective cells) as labels. We trained a K-Nearest Neighbor (KNN) clustering model on a random 20% subset of the total data and evaluated the decoding accuracy on the remaining 80%. This process was repeated 10,000 times, and the average decoding accuracy was calculated. This value was used as a quantitative measure of the quality of the clustering based on the ratio of reactivated neural ensemble types during SWRs.

### Statistical analysis

Both behavioral and neural data were analyzed using a statistical significance of α = 0.05. The one-sample Wilcoxon signed rank test was used only for testing the significance of behavioral performance (75% correct choices); for all other behavioral data, the significance of differences was tested using a two-sided Wilcoxon rank-sum test. Proportional differences between cell types (i.e., SF vs. MF) and SWR types were tested using the chi-squared test. The effects of region and cell type on cell firing activity, selectivity indices, or SWR participation rates were assessed by two-way analysis of variance (ANOVA), using an unpaired two-sample t-test with Bonferroni correction as a post hoc test.

## Acknowledgments Funding

This research was supported by National Research Foundation of Korea Grants 2019R1A2C2088799, 2021R1A4A2001803, 2022M3E5E8017723, and 2022R1I1A1A01069756.

## Author contributions

Conceptualization: JMS, SML, IL

Data curation: SML, IL

Formal Analysis: JMS, IL

Funding acquisition: IL

Methodology: SML, IL

Investigation: SML, IL

Project administration: IL

Resources: IL

Software: IL

Supervision: IL

Validation: IL

Visualization: JMS, SML, IL

Writing—original draft: SML, JMS, IL

Writing—review & editing: SML, JMS, IL

## Data and materials Availability

Customized Matlab codes were used for neural data analysis in the current study. The data and the code that support the findings of the present study are available from the corresponding author (I.L.) upon reasonable request. All data needed to evaluate the conclusions in the paper are present in the paper and/or the Supplementary Materials.

---

**Fig. S1.**
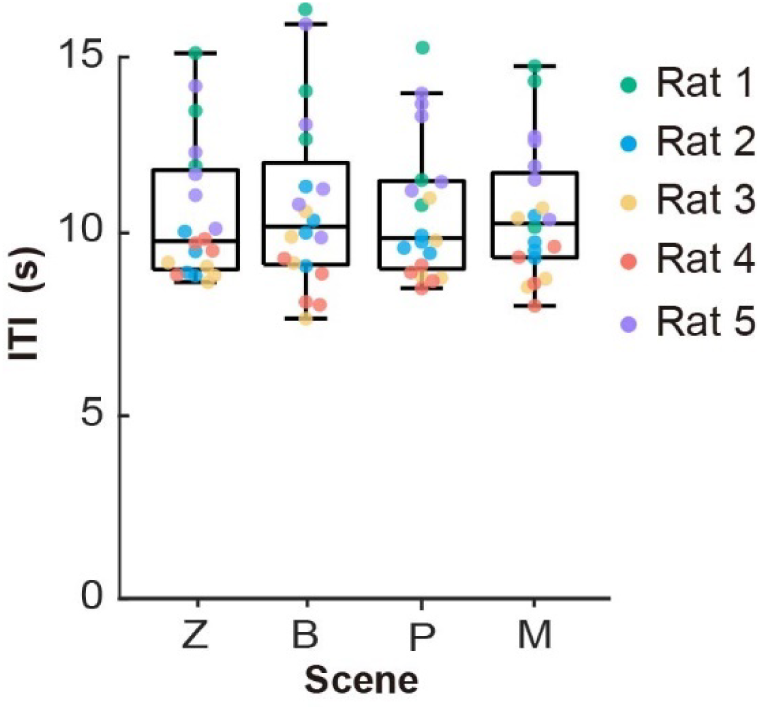
No difference in ITI among visual scenes. Comparison of ITIs among trial scene conditions. Box plot indicates median and interquartile values of the ITI for all rats in each scene. Each dot indicates the average ITI (excluding ITIs after incorrect trials) for each session, with sessions from the same rat depicted by the same color. Z, zebra; B, bamboo; P, pebbles; M, mountain.

**Fig. S2.**
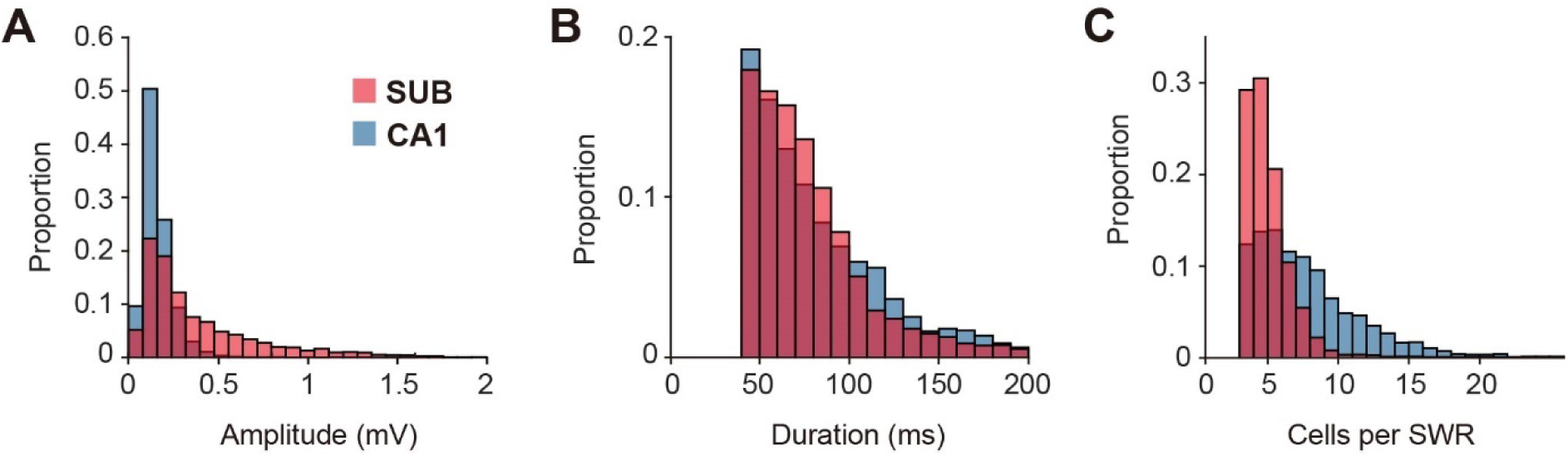
Differences in SWR properties between the subiculum and CA1. **(A)** Distribution of average filtered LFP amplitudes during SWRs (150-250 Hz). **(B)** Distribution of SWR durations. **(C)** Histogram of the number of activated cells per SWR. SUB, subiculum.

**Fig. S3.**
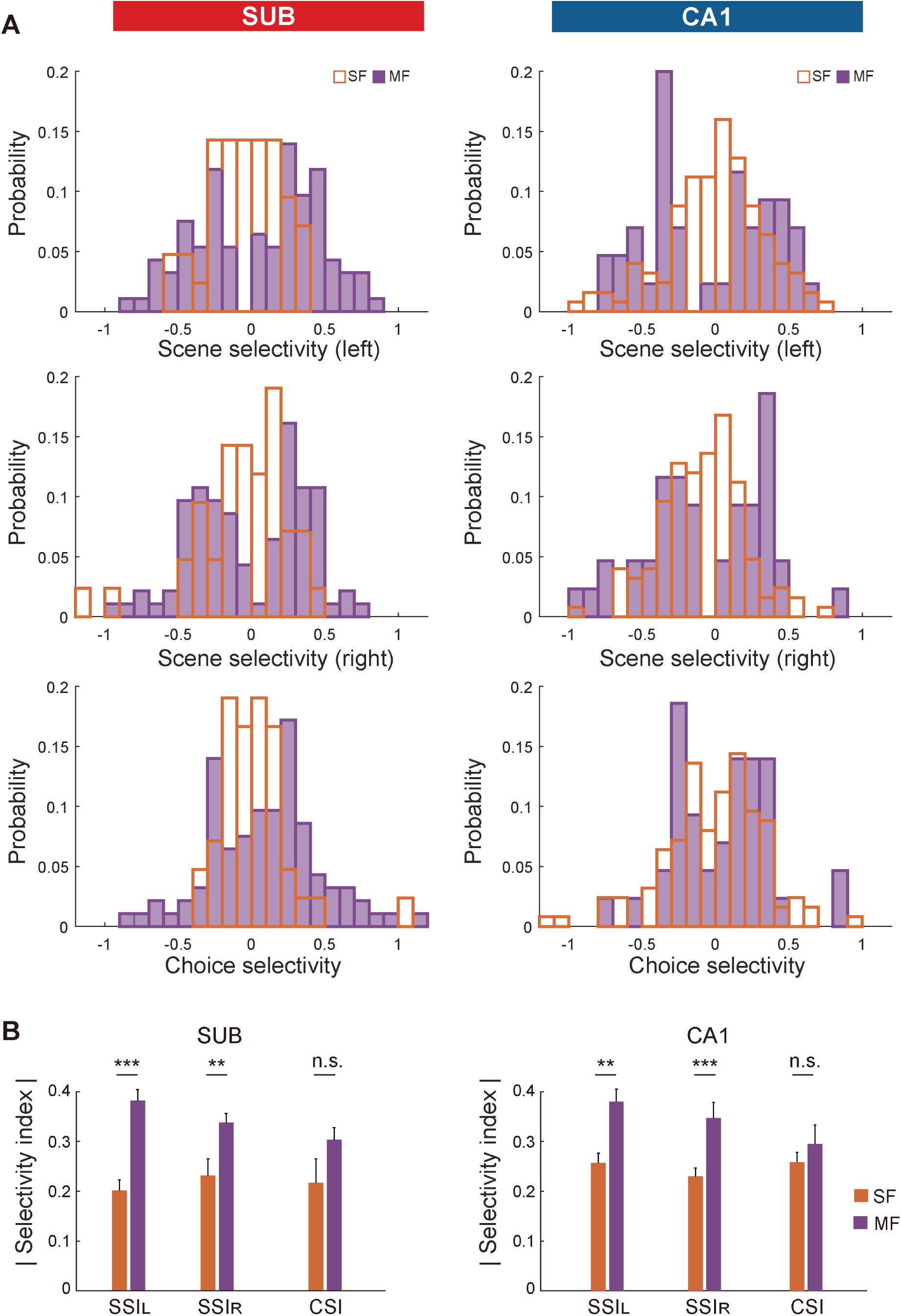
Selectivity index distribution of θ-phase–based subfields. **(A)** Histograms show selectivity index distribution of θ-phase–based subfields in the subiculum (left) and CA1 (right) for SSI_L_ (top) SSI_R_ (middle) and CSI (bottom). **(B)** Comparison of the absolute values of selectivity indices between SF and MF cells in the subiculum (left) and CA1 (right). Data are presented as means ± SEM. *** *p* < 0.001.

**Fig. S4.**
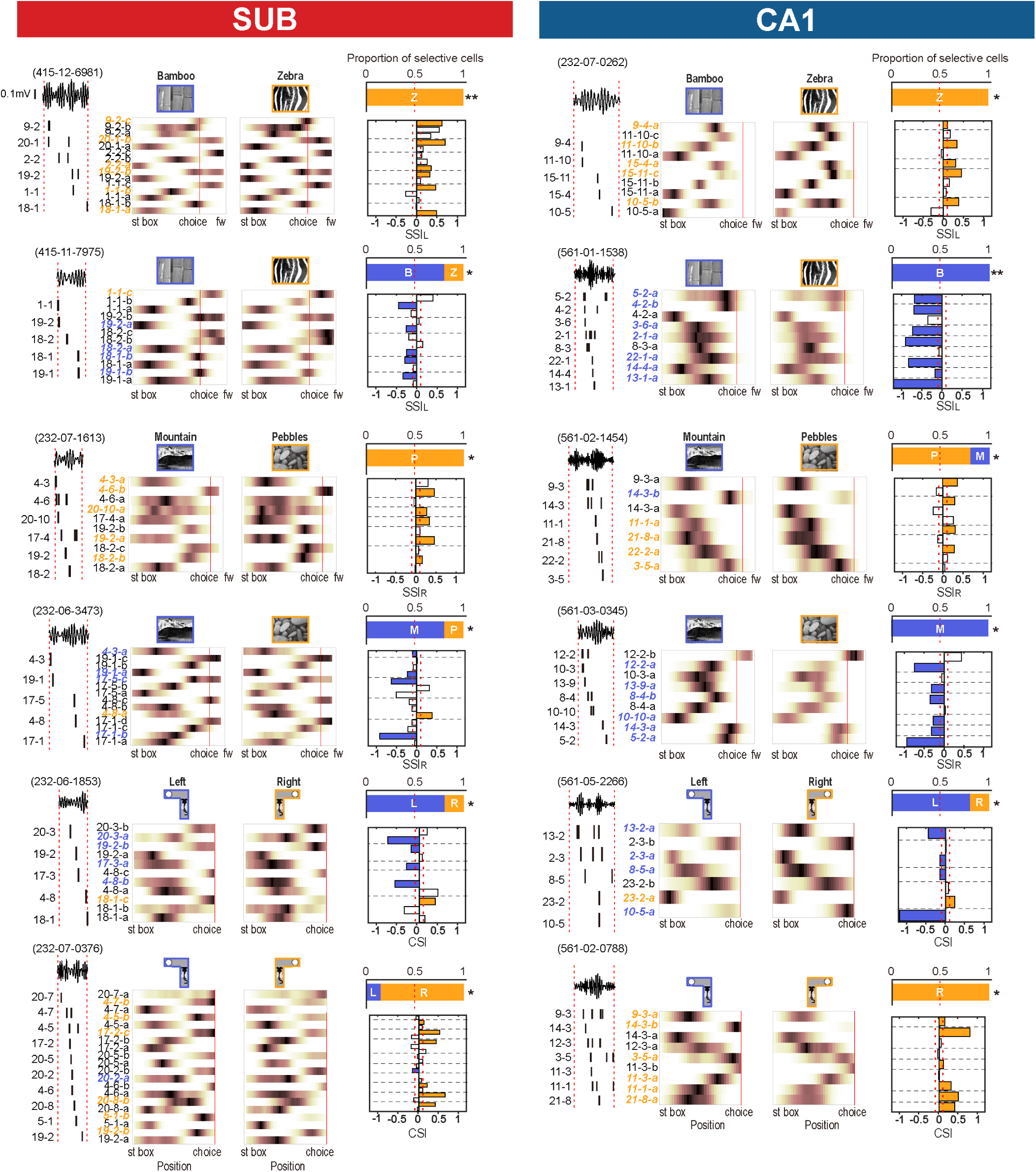
Original version of SWR examples in Fig. 4 that includes all θ-phase–based subfields of reactivated cells. Example SWRs with TDA reactivations in the subiculum (left) and CA1 (right). In each example, the leftmost part shows a filtered LFP trace and spiking activity of a cell ensemble within SWR boundaries. The middle colormaps represent the on-track firing rate maps of all θ-phase–based subfields for the selected task condition. For example, SWRs with choice reactivations, the ‘arm’ section of T-maze (from choice point to food well) is excluded from display since those sections are not used to calculate the choice selectivity index. The top right graph depicts the proportion of cells in the reactivated ensemble that are selective for certain trial conditions. The statistical significance of ensemble-level selectivity bias, calculated with a binomial test, is indicated as asterisks. The bottom right bar graphs are selectivity indices for θ-phase–based subfields, color-coded by trial condition. θ-phase–based subfields in the same cell are indicated by dotted lines.

**Fig. S5.**
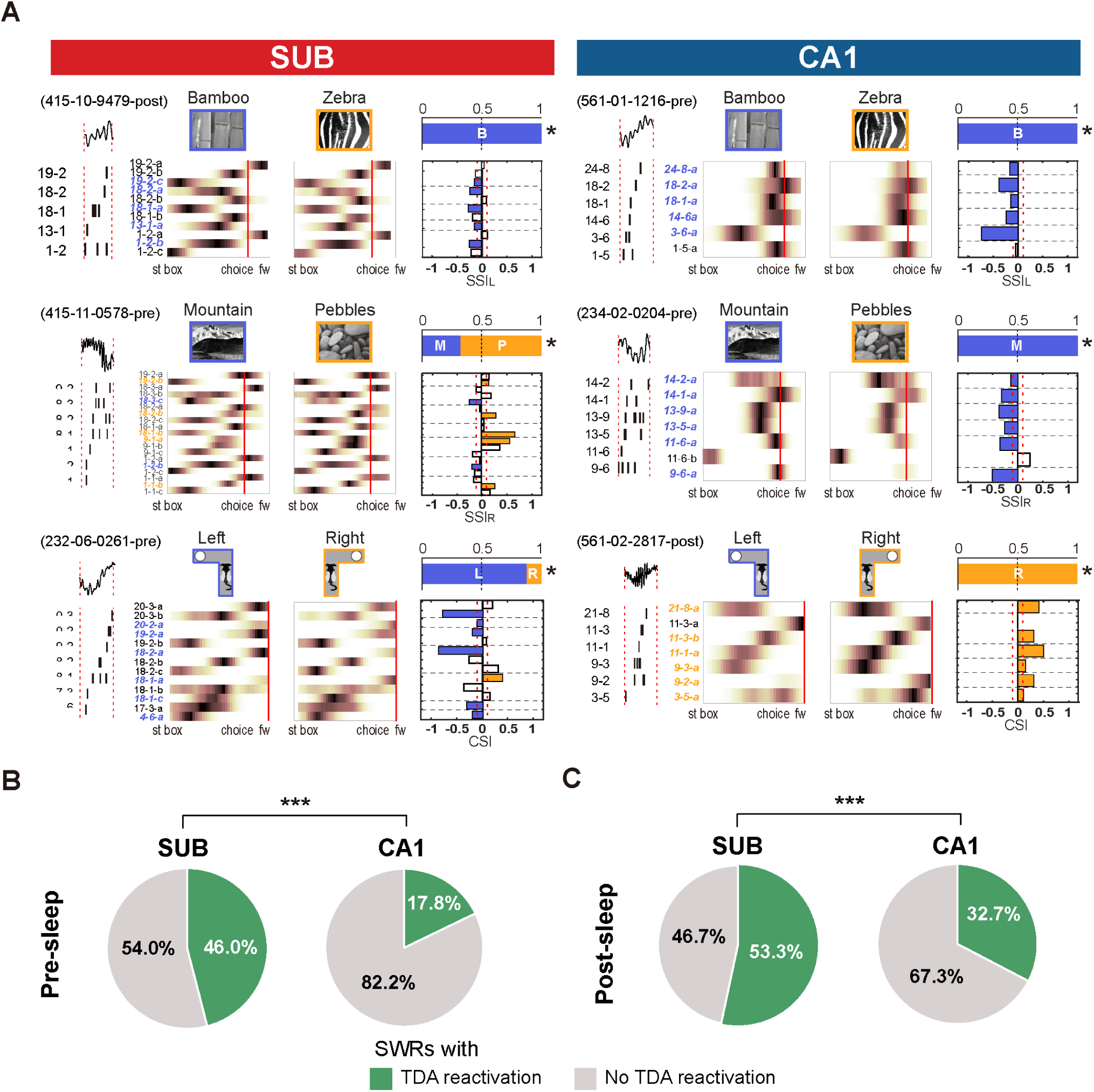
TDA reactivations during pre- and post-sleep period. **(A)** Example SWRs with TDA reactivations in the subiculum (left) and CA1 (right) during pre-sleep and post-sleep period. Each example is plotted in the same format as **Fig. S4**. **(B-C)** Proportion of TDA reactivations among all SWRs in the subiculum and CA1, during pre-sleep **(B)** and post-sleep **(C)** period. \**p* < 0.05, \*\*\**p* < 0.001. SUB, subiculum.

**Fig. S6.**
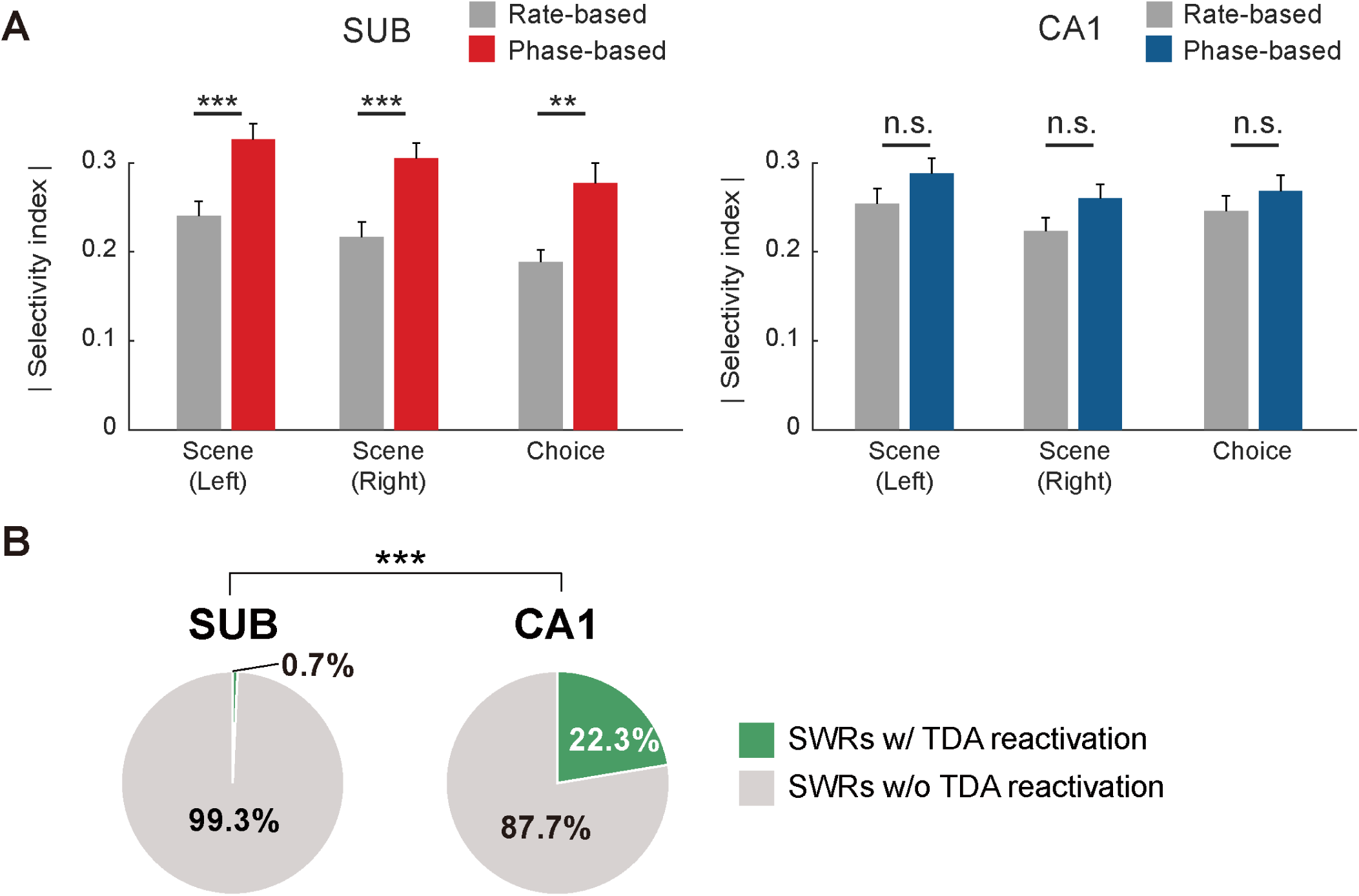
Reduced frequency of TDA reactivations after firing-rate–based field identification. **(A)** Bar graphs comparing the absolute values of the selectivity index calculated from rate-based field versus θ-phase–based field in the subiculum (left) and CA1 (right). In each region, the selectivity index is compared for the left-choice scene pair (zebra-bamboo), the right-choice scene pair (pebbles-mountain), and the choice pair. Data are presented as means ± SEM. **(B)** Proportion of TDA reactivations among all SWRs in the subiculum and CA1, when the selectivity index of each cell is calculated from the rate-based field. \*\**p* < 0.01, \*\*\**p* < 0.001. SUB, subiculum.

**Fig. S7.**
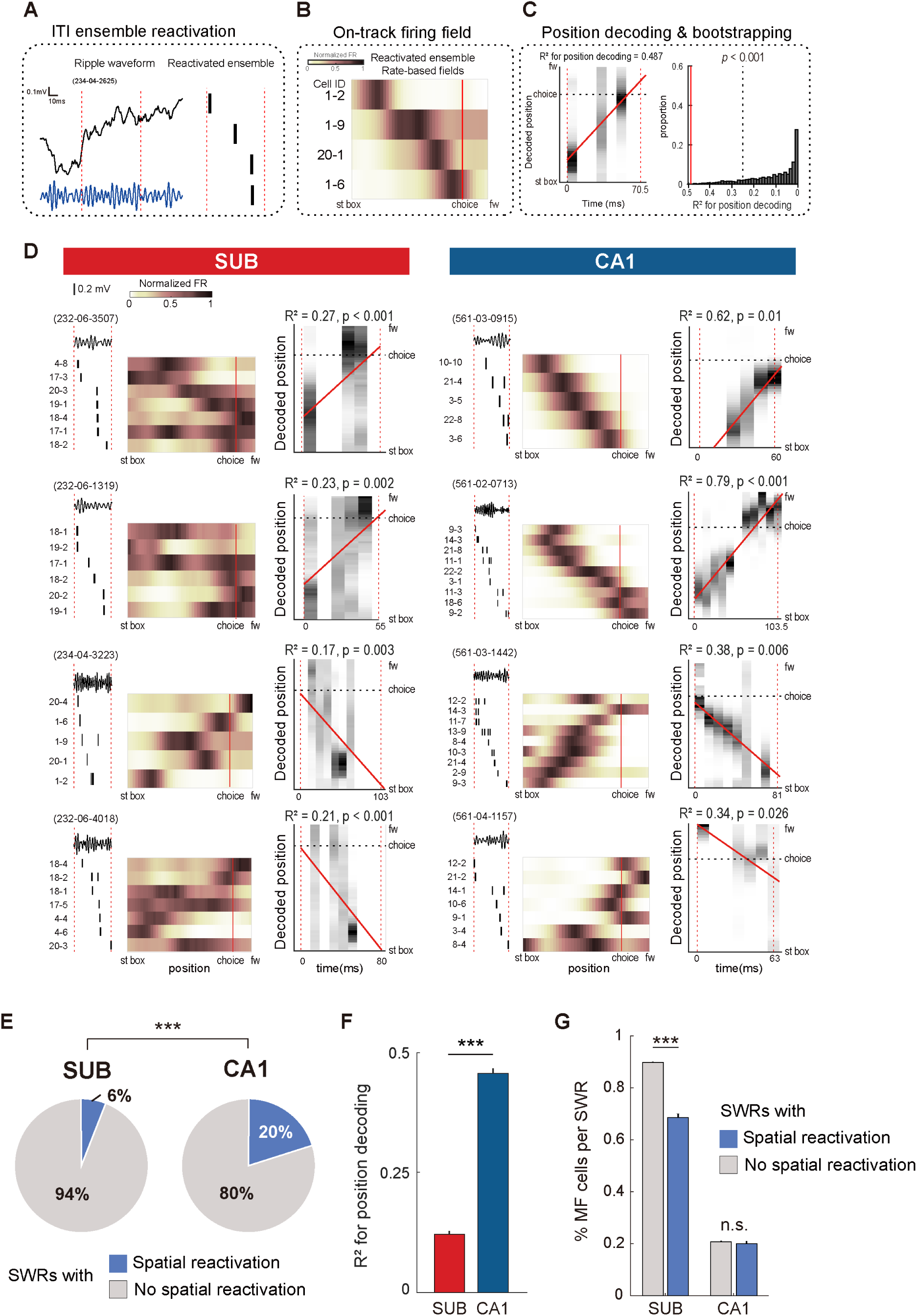
Spatial reactivation in SWRs is more prevalent in CA1 than in the subiculum. **(A)** An example SWR during an ITI is shown (depicted as in Fig. 3A). **(B)** On-track firing rate maps for each cell on the track. Each cell in a row is the cell in the same relative row as in the raster plot in (A). The numbers on the left side of each row indicate the cell ID. Firing rates in each map are normalized to the cell’s maximum firing rate (color bar at upper left). **(C)** Left panel: Posterior probability of the decoded position during the SWR. Red dashed line represents the linear regression line. The R^2^ value for the line is indicated at the top of the graph. Right panel: Bootstrap process used to calculate the significance of the regression line for the decoded position. Red line indicates the position of the example SWR’s R^2^ value from the distribution. The *p*-value represents the proportion of random distribution values larger than the example SWR’s R^2^ value. **(D)** Example SWRs with spatial reactivations in the subiculum (left panel) and CA1 (right panel). In each SWR, the leftmost line plot and raster plot show LFP changes and single-cell firing during the SWR. The middle colormaps show each single cell’s on-track firing rate map. The rightmost plot shows the posterior probability of the decoded position during each SWR (see Materials and Methods for detailed description). **(E)** Proportion of SWRs with spatial reactivations in the subiculum and CA1. **(F)** Comparison of R^2^ value (linear regression strength) for the posterior probability of the decoded position during SWRs between the subiculum and CA1. Data are presented as means ± SEM. **(G)** Proportional differences of MF cells calculated for each SWR event between SWRs with or without spatial reactivation. Data are presented as means ± SEM. \*\*\**p* < 0.001. St box, start box; choice, choice point; fw, food well; SUB, subiculum.

