## Supplementary figures and images for "Differential Reactivation of Task-Demand-Associated Firing Patterns in Subicular and CA1 Place Cells during a Hippocampal Memory Task"

### Fig.S1

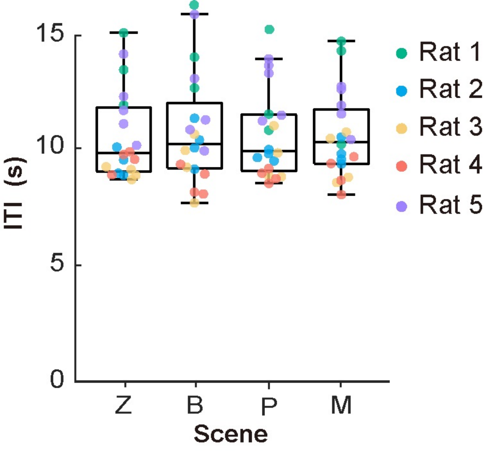

### Fig.S2

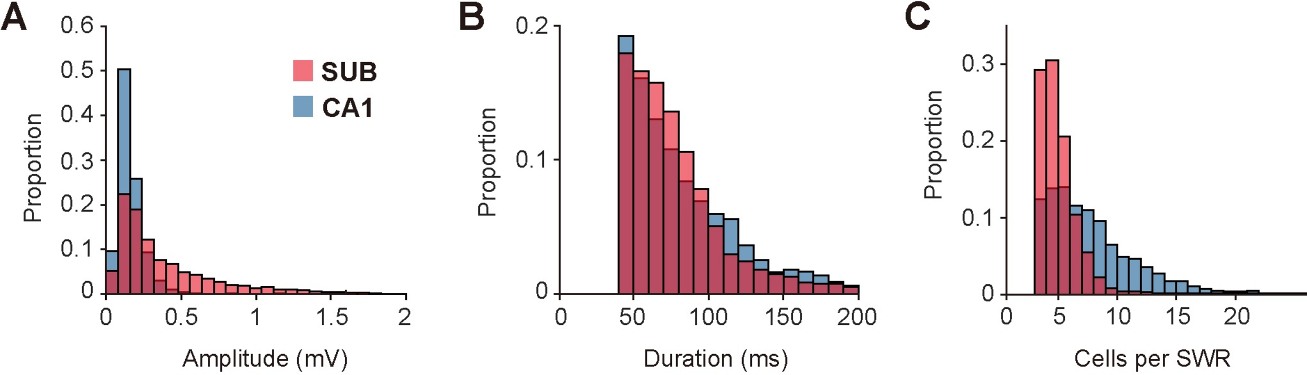

### Fig.S3

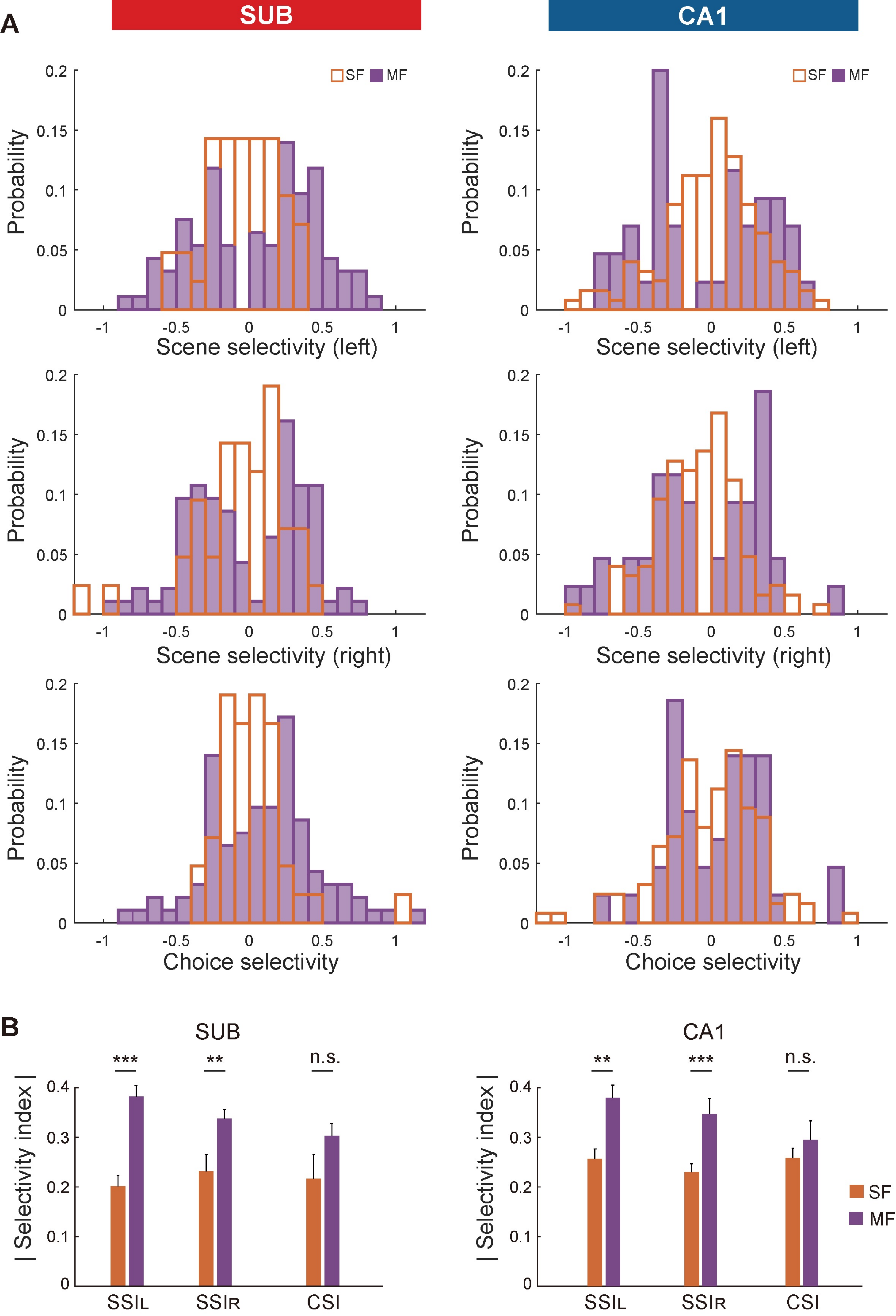

### Fig.S4

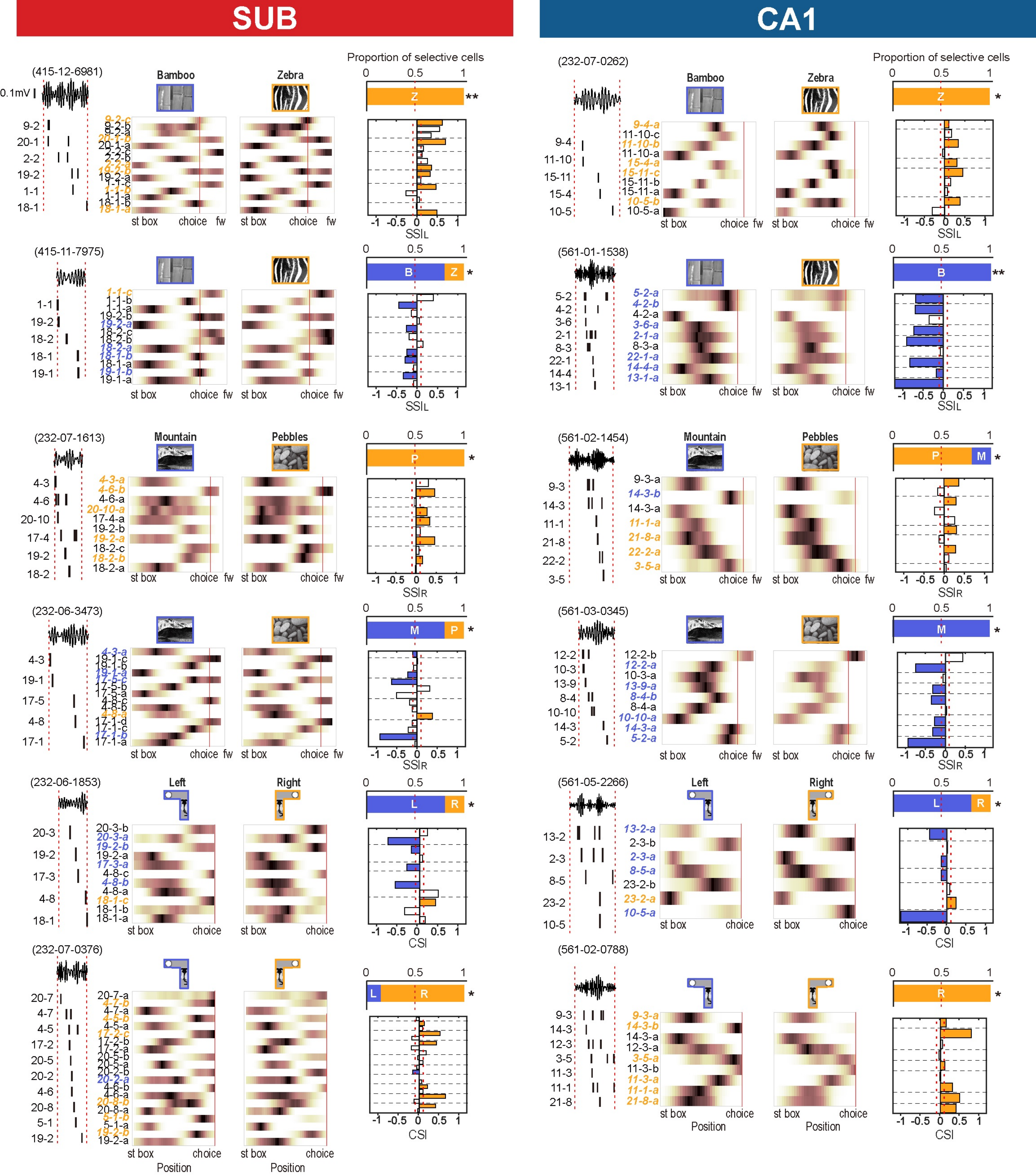

### Fig.S5

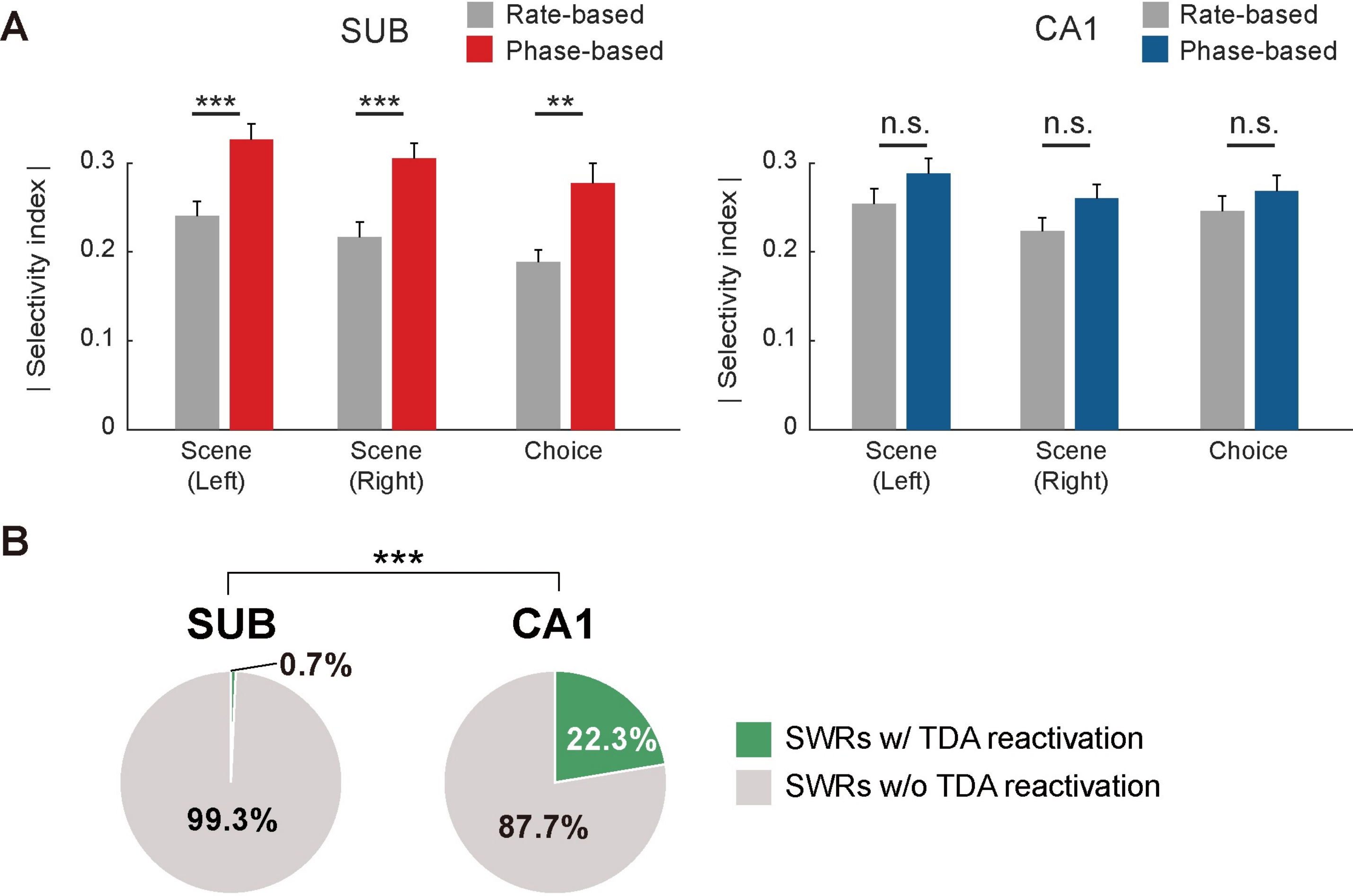

### Fig.S6

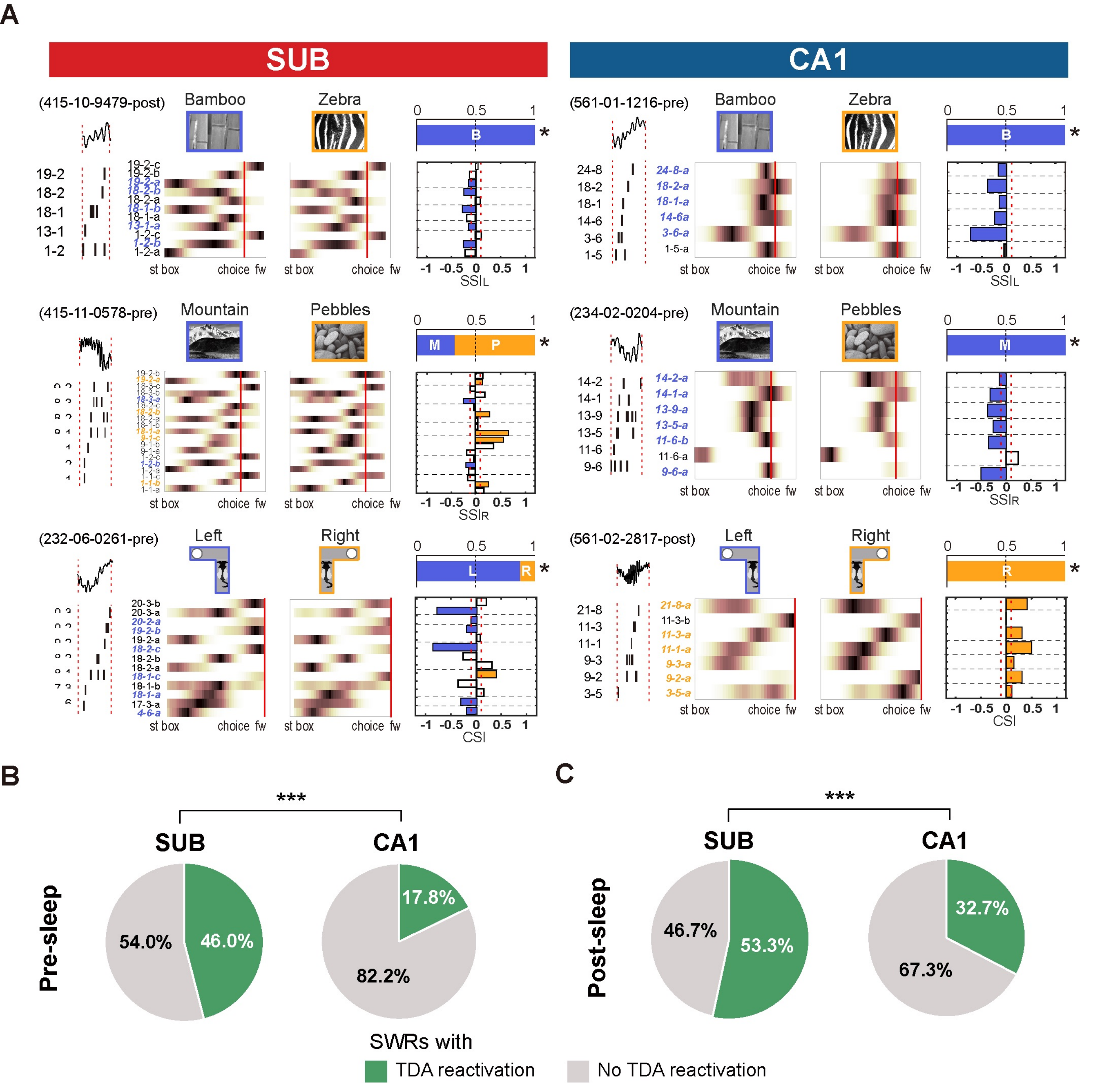

### Fig.S7

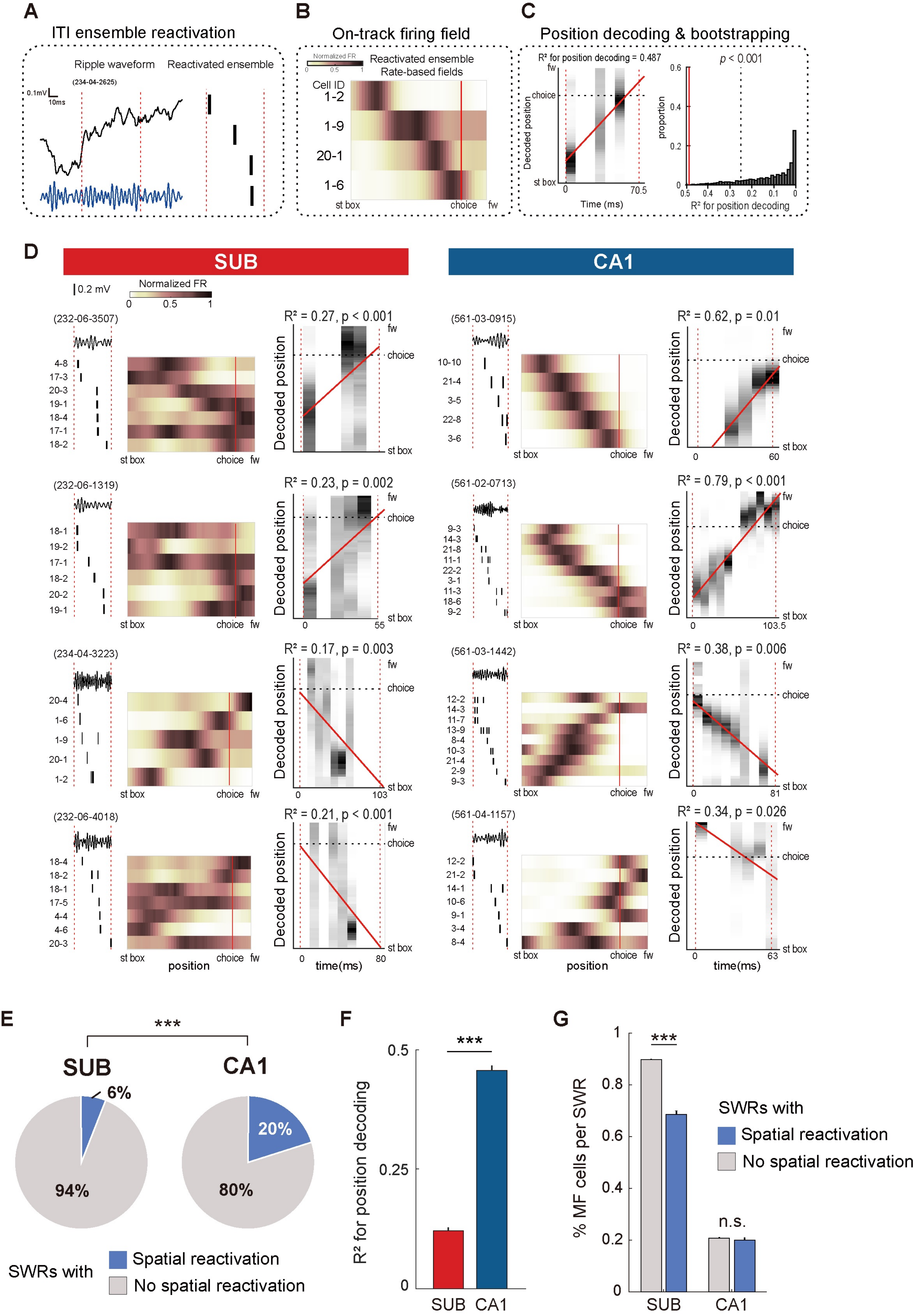
